# Meta-analysis of Drought-tolerant Genotypes in *Oryza sativa*: A Network-based Approach

**DOI:** 10.1101/450205

**Authors:** Sanchari Sircar, Nita Parekh

## Abstract

**Background:** Drought is a severe environmental stress. It is estimated that about 50% of the world rice production is affected mainly by drought. Apart from conventional breeding strategies to develop drought-tolerant crops, innovative computational approaches may provide insights into the underlying molecular mechanisms of stress response and identify drought-responsive markers. Here we propose a network-based computational approach involving a meta-analytic study of seven drought-tolerant rice genotypes under drought stress.

**Results:** Co-expression networks enable large-scale analysis of gene-pair associations and tightly coupled clusters that may represent coordinated biological processes. Considering differentially expressed genes in the co-expressed modules and supplementing external information such as, resistance/tolerance QTLs, transcription factors, network-based topological measures, we identify and prioritize drought-adaptive co-expressed gene modules and potential candidate genes. Using the candidate genes that are well-represented across the datasets as ‘seed’ genes, two drought-specific protein-protein interaction networks (PPINs) are constructed with up-and down-regulated genes. Cluster analysis of the up-regulated PPIN revealed ABA signaling pathway as a central process in drought response with a probable crosstalk with energy metabolic processes. Tightly coupled gene clusters representing up-regulation of core cellular respiratory processes and enhanced degradation of branched chain amino acids and cell wall metabolism are identified. Cluster analysis of down-regulated PPIN provides a snapshot of major processes associated with photosynthesis, growth, development and protein synthesis, most of which are shut down during drought. Differential regulation of phytohormones, e.g., jasmonic acid, cell wall metabolism, signaling and posttranslational modifications associated with biotic stress are elucidated. Functional characterization of topologically important, drought-responsive uncharacterized genes that may play a role in important processes such as ABA signaling, calcium signaling, photosynthesis and cell wall metabolism is discussed. Further transgenic studies on these genes may help in elucidating their biological role under stress conditions.

**Conclusion:** Currently, a large number of resources for rice functional genomics exist which are mostly underutilized by the scientific community. In this study, a computational approach integrating information from various resources such as gene co-expression networks, protein-protein interactions and pathway-level information is proposed to provide a systems-level view of complex drought-responsive processes across the drought-tolerant genotypes.

## Background

Drought is an inevitable consequence of drastic climate change. A look at the World Drought Map for the year 2017 from the Global Drought Information System [1] reveals the severity and intensity of drought across the globe. At this juncture, meeting the nutritional demands for an ever-increasing population is a challenge for policy makers. Although drought and heat waves have destroyed nearly a tenth of the cereal harvests, populations across the world continue to depend on them for food.

Irrigated rice occupies half the world’s rice field and accounts for ∼75% of the global rice supply. This makes its harvest and yield highly susceptible to climate change. Abiotic stress conditions like drought induces a wide array of physiological, morphological and molecular changes which might provide cues towards drought-responsive/adaptive mechanisms. However, these approaches are limited by the fact that nearly ∼50% of the rice genes lack functional annotation for biological processes [2]. High-throughput technologies together with computational methods can be effective in functional characterization of rice genes, reducing the search space for identification of novel candidate genes and elucidating biological processes under various environmental conditions. Various drought tolerant rice genotypes exist, such as upland *indica* variety Bala with a high stomatal sensitivity to water loss and upland *japonica* variety Azucena with thick and deep roots. The deep-rooted and stress tolerant *aus* rice cultivar Nagina 22 (N22) is known to have enhanced enzymatic activity, α-linolenic acid metabolism and heat shock proteins [3,4]. Studies performed with indigenous cultivar Dagad Deshi [5] also identified quantitative traits associated with grain yield. The drought tolerant upland variety IRAT109 is known for its superior yield and other parameters compared to the lowland Zhenshan97 under drought stress [6] and have been used for studying traits like osmotic potential [7]. Thus there has been a continuous effort in conventional breeding strategies to develop high-yielding drought-tolerant rice varieties.

Currently, a large number of gene-expression data corresponding to drought tolerant genotypes are available in public domain. Systematic meta-analysis of these datasets can help in increasing the reproducibility and provide new biological insights. Numerous methods are available to merge gene expression data and remove batch effects, *viz*., Z-score standardization, mean centering [8], cross-platform normalization [9], empirical Bayes method for smaller batch size [10], etc. Network-based approaches help in summarizing large-scale data with thousands of genes into manageable clusters or functional modules. Further, these approaches help to prioritize important genes based on their topological parameters such as degree, betweenness and eigen-gene centralities, apart from fold-change and functional enrichment. For example, co-expression networks are popularly used to draw relationship between genes and identify tightly coordinated biological processes and functional characterization of novel genes [11–15]. An integrative approach combining co-expression networks with protein-protein interactions and metabolic pathway information can help in associating genotype to phenotype to a considerable extent [16–18].

In this study, we carried out meta-analysis of drought-tolerant rice genotypes under drought stress using network-based approach with the objective of identifying gene signatures and processes that are hallmark of drought tolerant species. For this analysis, from the six microarray datasets containing seven drought-tolerant genotypes (Azucena, Bala, Nagina, Dagad deshi, Vandana, IRAT109 and *Oryza* DK151), 57 samples corresponding to the shoot tissue and seedlings from whole plant (Affymetrix platform) are considered (**Table 1**). Samples corresponding to drought-sensitive genotypes or root tissue in these datasets are not considered for the analysis (see **Table S1, Additional File 1** for sample details). The R package, Weighted Gene Co-expression Network Analysis (WGCNA), is used to capture interaction patterns between genes [19] resulting in co-expressed gene modules. These co-expressed modules are associated with additional information such as enrichment of differentially expressed genes, transcription factors, known stress/tolerance associated QTLs, module-trait associations and gene ontology, to identify drought-responsive modules. The key driver genes in these modules are identified using topological network parameters and up-and down-regulated protein-protein interaction networks are constructed from known interaction of these genes. Tightly coupled gene clusters that may be associated to various signaling and metabolic processes likely to be affected due to drought are analyzed.

**Table 1.** Drought-tolerant samples from Affymetrix datasets from NCBI-GEO and ArrayExpress considered for the meta-analytic are listed.

| Microarray studies<br>(Total No. of Samples) | Title of the Experimental Studies | No. of Control + Drought Samples | Tissue Samples Considered |
| --- | --- | --- | --- |

|  |  | Considered |  |
| --- | --- | --- | --- |
| **GSE41647\***<br>(18) | Transcriptome profiling for drought tolerant and susceptible cultivars of Indica rice | 9<br>Dagad deshi<br>(DD) | Seedlings<br>(whole plant) |
| **E-MEXP-2401\***<br>(12) | Transcription profiling of *Oryza sativa* subtypes Cultivar Nagina-22 (N22) and IR64 subtypes under normal and drought conditions | 6<br>Nagina22<br>(N22) | Seedlings<br>(whole plant) |
| **GSE21651\***<br>(16) | Differential expression for salt and drought stress from tolerant and sensitive lines | 4<br>Vandana | Seedlings<br>(leaf) |
| **GSE26280<sup>#</sup>**<br>(36) | Genome-wide temporal-spatial gene expression profiling of drought responsiveness in rice | 6+6+6<br>*Oryza* DK151 | Leaves<br>(tillering + panicle<br>elongation + booting)<br>(vegetative +<br>reproductive stages) |
| **GSE24048**<br>(12) | Expression data from field droughted rice plants | 12<br>6 Bala + 6<br>Azucena | Leaves<br>(vegetative stage) |
| **GSE25176\***<br>(16) | Expression data from rice varieties IRAT109 (resistant) and ZS97 (sensitive) for drought stress treatment in flag leaves | 8<br>IRAT109 | Flag leaf<br>(reproductive stage) |

\*These datasets also have drought-sensitive samples which are not used in this study
<sup>#</sup>This dataset had root tissue samples from tillering and panicle elongation stages and young panicle tissue in booting stage which are not used in this study.

## Methods

### Data Pre-processing

A first step in any transcriptomic studies is to check the data quality. All the samples are found to pass quality check on using the package, ArrayQualityMetrics that assesses reproducibility, outlier arrays, batch effects and computes measures of signal-to-noise ratio [20]. Datasets are normalized using Robust Multiarray Averaging (RMA) package to remove systematic variations due to different experimental conditions, dye effects, uneven hybridizations, etc. [21]. In a meta-analytic approach, when multiple datasets from different experimental conditions are combined, batch-specific variations need to be eliminated. For this, COMBAT, an Empirical Bayes method in the R package inSilicoMerging, is used to merge the expression values across the datasets [10]. Probes having no gene annotations, mapping to more than one gene annotation (ambiguous probes), or having very low intensity values (< 20 across all samples) are removed. For multiple probes mapping to the same gene, the one exhibiting a higher coefficient of variation across the samples is considered. This resulted in 14,270 genes that are used for the construction of co-expression network.

### Co-expression Network Construction

The weighted gene correlation network analysis (WGCNA) package is used to construct a ‘signed’ co-expression network of drought tolerant rice samples from normalized, log2-transformed expression matrix across 14,270 genes and 56 samples (one sample, GSM645335 was detected as an outlier on performing sample-wise clustering in WGCNA and removed). The unsigned networks use absolute value of correlations, 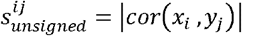, and are unable to distinguish between gene activation 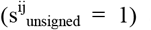 and gene repression 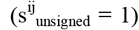, leading to loss of biological information [22]. Hence, here we construct a signed co-expression network, taking into account the ‘sign’ of correlation between expression profiles of genes and the similarity measure in this case is defined as:

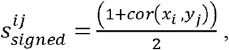

where *x_i_* and *x_j_* are the expression profiles of genes *i* and *j* across the microarray samples [22]. Here, 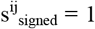 corresponds to positive correlation, 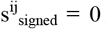, negative correlation and 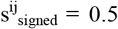, no correlation, thereby distinguishing between positively and negatively correlated genes. The similarity between gene-pairs is computed using signed Pearson’s correlation matrix, scaled to power β = 18 (approximates scale free-topology criterion), and the parameters of the signed co-expression constructed are summarized in **Table 2**. The function *block-wiseModules* is used for hierarchical clustering of genes using Dynamic Tree Cut approach [23] with maximum block size = 15000, minimum module size = 100, “cut height” = 0.995 and “deep split” = 2. This resulted in 13 co-expressed gene modules ranging in size from 2686 (turquoise) to 250 (salmon) genes, and 1431 genes are left unclustered (in grey module). The co-expression network with all the annotations are submitted as **Table S2, Additional File 1.**

**Table 2.** Parameters for the construction of signed, weighted gene co-expression network is summarized.

| Genotype | No. of Samples | No. of Genes | $\beta$ Cutoff | $R^2$ Scale-free fit | Mean k | Median k | Max k | No. of Modules |
| --- | --- | --- | --- | --- | --- | --- | --- | --- |
| Drought Tolerant | 56 | 14270 | 18 | 0.85 | 53.4 | 30.9 | 331 | 13 |

### Statistical and Biological Significance of Network Modules

#### Statistical Significance of Network Modules

To assess the statistical robustness of the co-expressed modules, module quality statistics are computed by re-sampling the dataset using the *modulePreservation* function in WGCNA. It randomly permutes the gene labels 200 times and computes various network quality statistics such as density, module membership, connectivity, etc. for each module. The log *p*-values and *Z*-scores for each module are summarized as *psummary* and *Zsummary* (see **Table S3, Additional File 1**). The *Z*-score provides evidence that a module is preserved more significantly than a random sampling of genes and *p*-value gives the probability of seeing the module quality statistic in a random sampling of genes of the same size. It is observed that the *psummary* value is very low (∼ 0.0) and *Zsummary* > 10, providing strong evidence of network connectivity preservation and robustness of the co-expressed modules [24].

#### Biological Relevance of Network Modules

Gene ontology and enrichment analysis of the modules carried out using agriGO [25] indicates probable pathways captured by the module-genes. After merging the six datasets using inSilicoMerging, differentially expressed genes (DEGs) are identified using the criteria of average fold-change across 6 datasets ≥ |1.2| and a *t*-test with *p*-value ≤ 0.05. A total of 6441 DEGs are identified and mapped on to the co-expressed modules (**Table 3**). We observe a clear demarcation in the distribution of up-and down-regulated genes on the modules, clearly showing the effect of constructing a ‘signed’ network. The Turquoise, Yellow, Tan and Brown modules are enriched with only up-regulated genes (3012) while Blue, Green, Red, Magenta, Salmon and Purple modules contain mainly down-regulated genes (3235) and these modules are likely to capture processes that are activated or repressed in response to drought, respectively.

**Table 3.** Co-expressed modules with percentage DEGs, transcription factors (TFs) and GO enriched terms are shown. Modules are ordered by their size.

| Module | Size | DEGs (%age) |  | TFs<br>(from PlnTFDB) |  | GO Analysis using agriGO |
| --- | --- | --- | --- | --- | --- | --- |
|  |  | Up | Down | Up | Down |  |
| Turquoise | 2686 | 2103<br>(78.3) | 0 | 171 | 0 | regulation of cellular process (204), catabolic process (63), transcription (165), localization (160) |
| Blue | 2500 | 0 | 1861<br>(74.4) | 0 | 99 | small molecule metabolic process (156), photosynthesis (36), cellular nitrogen compound metabolic process (60), oxoacid metabolic process (78), establishment of localization (179) |
| Brown | 1327 | 247<br>(18.6) | 0 | 7 | 0 | RNA processing (25), gene expression (119), DNA repair (17), lipoprotein metabolic process (7) |
| Yellow | 1203 | 561<br>(46.6) | 3<br>(0.25) | 43 | 0 | Transport (110), catabolic process (42), establishment of localization (104), small molecule metabolic process (70), generation of precursor metabolites and energy (31) |
| Green | 1192 | 0 | 713<br>(60) | 0 | 57 | protein modification process (121), post-translational protein modification (112), phosphate metabolic process (105) |
| Red | 884 | 9<br>(1) | 286<br>(32.3) | 1 | 4 | ncRNA metabolic process (43), RNA processing (41), translation (67), gene expression (124), RNA modification (19), ribosome biogenesis (20) |
| Black | 677 | 10<br>(1.5) | 20<br>(3) | 2 | 4 | protein modification process (73), transport (62), macromolecule modification (73), signal transduction (21) |
| Pink | 633 | 42<br>(6.6) | 22<br>(3.5) | 0 | 1 | establishment of localization in cell (38), intracellular transport (34), vesicle-mediated transport (30) |
| Magenta | 415 | 0 | 187<br>(45) | 0 | 8 | cellular glucan metabolic process (10), polysaccharide metabolic process (11), cellulose biosynthetic process (6) |
| Purple | 390 | 0 | 95<br>(24.4) | 0 | 6 | cellular protein metabolic process (48) |
| GreenYellow<br>w | 347 | 2 (0.6) | 86<br>(24.8) | 0 | 3 | Translation (75), gene expression (92), cellular protein metabolic process (85), cellular biosynthetic process (103) |
| Tan | 335 | 101<br>(30) | 0 | 8 | 0 | - |
| Salmon | 250 | 0 | 93<br>(37.2) | 0 | 9 | gene expression (35), regulation of metabolic process (24), regulation of transcription (22) |

### Construction of Minimal Drought-adaptive PPI Networks

The modules obtained in the co-expression network are in general quite large and as such are not very useful. Also, fold-change alone does not say much about the systems-level significance of a gene. To address some of these issues, we additionally consider network topological measures such as intra-modular connectivity/degree centrality (k_IM_), which help in identifying ‘hub’ genes and eigengene-based centrality (k_ME_), which is a measure of module membership of a gene obtained by correlating its expression profile with eigengene of the corresponding module, to filter important stress-responsive genes. Top 20% genes ranked based on high k_IM_ and k_ME_ values that are also differentially expressed (fold-change ≥ |1.2|) are selected from each of the 4 up-regulated (Turquoise, Yellow, Tan and Brown) and 6 down-regulated (Blue, Green, Red, Magenta, Purple and Salmon) drought-responsive modules. This resulted in 754 up-regulated and 893 down-regulated genes. These stress-responsive genes are used as ‘seeds’ to search the database of protein-protein interaction networks, STRING [26] and a total of 939 interactions between 425 up-regulated genes and 8639 interactions between 640 down-regulated genes are identified. The remaining unmapped genes are queried in co-function networks, RiceNet [27] and their orthologs in AraNet [28], and additional interactions (between 76 up-regulated and 80 down-regulated genes respectively) that are conserved between both RiceNet and AraNet are retrieved. This resulted in two PPINs which we refer to as the up-regulated drought tolerant network (uDTN) consisting of 466 nodes and 1015 edges, and the down-regulated drought tolerant network (dDTN), consisting of 665 nodes and 8719 edges. The largest connected component in these two PPINs is considered for further analysis. The largest connected component of uDTN containing 422 nodes and 990 edges, and that of dDTN with 623 nodes and 8627 edges, is observed to follow an approximate power law degree distribution (with R^2^ value of 0.85 and 0.76 respectively), indicating scale-invariant topology. These two components are then subjected to Markov Cluster Algorithm (MCL). Tightly coupled gene clusters (13 in uDTN and 14 in dDTN) of size ≥ 5 and showing significant functional enrichment in terms of pathway, GO or domain in STRING database are identified. We expect these gene clusters to provide insight into the biological processes that are activated or repressed in response to drought stress in drought tolerant varieties. The networks are visualized using Cytoscape [29] and topological analysis is carried out using NetworkAnalyzer plugin.

## Results

### Identification of Drought-adaptive Modules

The signed weighted co-expression network of 14,270 genes resulted in 13 co-expressed modules. These modules are large, comprising hundreds and thousands of genes. To identify key drought-responsive processes we carry out various statistical operations discussed below.

### Differential Gene Expression Analysis and GO Enrichment

GO enrichment analysis of the 13 co-expressed gene modules is carried out using agriGO [25] and the top enriched GO terms are given in **Table 3**. The number of up-and down-regulated genes (DEGs) and known rice transcription factors (TFs) extracted from PlnTFDB [30] are identified and mapped on these modules. The results are summarized in **Table 3.** The DEGs are identified based on fold-change (≥ |1.2|) and filtered using *t*-test with *p*-value (≤ 0.05). It is evident from **Table 3** that Turquoise module has the highest number of up-regulated genes (2103). It harbors genes involved in the regulation of various cellular processes and transcription factors such as bZIPs, NACs and MYBs, which are known to be involved in drought stress. The next major up-regulated module is the Yellow module with 561 DEGs. Apart from being involved in transport and localization, this module also consists of genes involved in catabolic processes and energy generating processes. Blue module has a large number of down-regulated genes (1861), majority of which are involved in photosynthesis and associated metabolic processes known to be down-regulated in shoot tissue during drought [31,32]. Green and Magenta modules also contain a significant fraction of down-regulated genes (713 and 187 respectively) and are involved in protein synthesis and cellulose metabolism respectively. Both these processes are known to be hampered during drought as a result of growth retardation and damage to membranes [33]. The Red module contains 286 down-regulated genes involved in gene expression, ribosome biogenesis and RNA processing, indicating a finer regulatory mechanism controlling protein synthesis. Based on GO enrichment analysis and distribution of DEGs, we associate 4 modules (Turquoise, Yellow, Brown and Tan) with up-regulated processes and 6 modules (Blue, Green, Red, Magenta, Purple and Salmon) with down-regulated processes.

Transcription factors play an important role in translating stress-induced signals to cellular responses by binding to specific *cis*-elements of downstream target genes. A number of regulons, *viz.*, like AREB/ABF regulon, DREB1/DREB2 regulon, NAC regulon, MYB and WRKY TFs have been identified to play an important role in stress response. Interactions between the regulons such as AREB/ABFs and NACs have been reported where NAC TFs regulate ABA biosynthetic genes. NAC-induced regulation of ABA regulons and presence of ABRE elements in the promoter regions of SNAC TFs hints at complex cross talks between the regulons [34–36]. Similarly, MYB TFs have been reported to be involved in stomatal patterning, regulation of guard cells controlling stomatal movements indicating an overlap with the ABA signals [37–39]. The DREB genes have been shown to physically interact with AREB/ABFs [40]. We show that our co-expressed modules are able to capture this inter-dependence of TFs and genes shown to be involved in drought tolerance.

To identify the differentially expressed TFs associated with the regulons, we analyzed their distribution across the 13 co-expressed modules. For this analysis, known rice transcription factors (2385 genes) are downloaded from PlnTFDB (3.0) [30]. Of the 776 TFs that mapped to the co-expression network, 232 are up-regulated and 192 down-regulated. The distribution of these differentially expressed TFs on the co-expressed modules is summarized in **Table 3**. We observe that 171 of the up-regulated TFs mapped to Turquoise, 43 to Yellow, and 8 and 7 to Tan and Brown modules respectively, while 99 of the down-regulated TFs mapped to Blue, 57 to Green, 9 to Salmon, 8 to Magenta, etc. In **Figure 1(A)** and **1(B)**, the total number of known members of various TF families that mapped to our co-expression network is depicted in ‘black’, while the ‘red’ and ‘blue’ bars depict the number of known up-and down-regulated TFs respectively. It is evident from **Figure 1(A)** that high numbers of up-regulated TFs belong to bZIP family followed by NAC, MYB-related and AP2-EREBP TFs, while from **Figure 1(B)** we observe that high numbers of HB, Orphans and bHLH are down-regulated. In plants, bZIP TFs regulate diverse processes including seed germination, flowering, photomorphogenesis, abiotic stress and ABA signaling [41]. Among the 21 up-regulated bZIP TFs, 17 are mapped to Turquoise module, including the well characterized OsABF1 (OsABI5/OREB1/OsbZIP10) TF. Recent study indicates that OsABF1 is a universal positive regulator of drought tolerance and ABA signaling in rice. It directly regulates OsbZIP23, OsbZIP46, and OsbZIP72 all of which are a part of Turquoise module [42]. It is also noted that both OsABF1 and OsbZIP23 are ‘hubs’ in the module (top 20% high degree genes). The NAC regulon plays an important role in stress response under various abiotic and biotic stress conditions as well as in various developmental programs such as lateral root development, flower development, formation of secondary walls, maintenance of secondary walls, etc. [43]. Among the 19 up-regulated NAC TFs, 14 mapped to Turquoise module, 4 to Yellow module and 1 to Brown module. For example, OsNAC52 which has been shown to be involved in drought tolerance [44] and LOC_Os07g48450 (yet to be characterized in drought stress) are observed to have a significant fold-change and high degree in the Turquoise module. The MYB-type TFs are one of the most abundant TFs in plants and play essential roles in development, growth and stress response and have been classified based on the number of MYB repeats [45]. We observe 18 MYB-related and 9 MYB TFs that are up-regulated, many of which are not yet functionally characterized for stress tolerance except for OsMYB55, reported to be involved in heat tolerance and amino acid metabolism [46] and OsMYB91, in plant growth and salt tolerance [46]. We also other 8 MYB and 10 MYB-related TFs to be down-regulated (**Figure 1(B)**). Among these, the MYB TF LOC_Os01g44390 (OsRAD1), involved in floral development, is significantly down-regulated and is also a ‘hub’ gene in the Blue module. The other class of important TFs are AP2-EREBPs which are highly abundant and involved in development, sugar signaling, ethylene response, pathogen response and abiotic stress [47,48]. We observe 17 up-regulated AP2-EREBPs TFs in our co-expression network, of which 13 map to Turquoise module and 2 each to Yellow and Tan modules. The DREB genes of this TF family bind to dehydration responsive element (DRE)/C-repeat (CRT) regions and activate LEAs, dehydrins, starch-degrading enzymes, Hsps, etc. [49] and play a role in stress response pathway. In **Figure 1 (B)**, we observe 17 homeobox genes to be down-regulated. These genes encode TFs which are important regulators for development, cell fate and body plan specification [46]. Majority of these TFs are part of the Blue module. Transcription factors (9 up-regulated and 14 down-regulated) belong to the Orphans category in PlnTFDB, which corresponds to TFs that could not be categorized into existing TF families. We observe 4 WRKY TFs in Green module, *viz*., WRKY45 (associated with disease resistance [50]), WRKY47 (positive regulator drought in *P_SARK_::IPT* plants [51]), WRKY68 and WRKY74, and WRKY69, WRKY109 part of Black and Blue module respectively. All these WRKYs are down-regulated.

**Figure 1:**
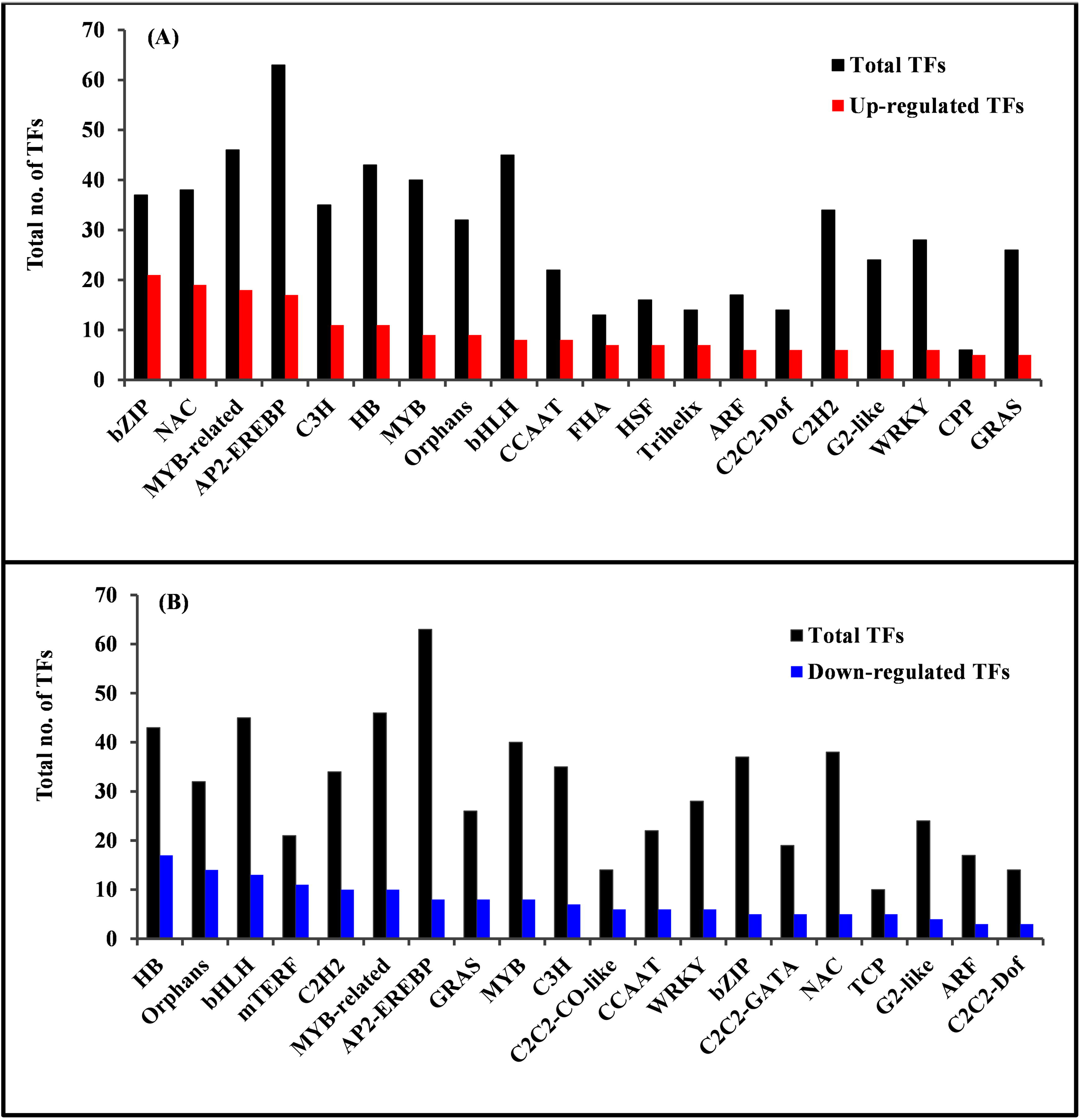
Transcription factor (TF) families identified in our co-expression network are ranked based on number of differentially expressed members (fold-change ≥ |1.2| and p-value ≤ 0.05). In (A) top 20 up-regulated TF families (‘red’ bars), and (B) top 20 down-regulated TF families (‘blue’ bars) are shown. The vertical bars in ‘black’ depict the total number of members of the respective TF families in (A) and (B). The gene-ID-TF mapping is taken from PlnTFDB v3.0.

Thus, we observe that the coordinated up-regulation of TFs corresponding to different regulons are part of the same modules (mainly Turquoise and Yellow), while TFs associated with disease resistance, flowering and development are part of the down-regulated (mainly Blue and Green) modules.

### QTL Genes Mapping to Co-expressed Modules

To further validate the biological significance of the 10 co-expressed modules in response to drought stress, we mapped known quantitative trait locus (QTL) genes on these modules. For this analysis, genes spanning QTLs that are associated with resistance/tolerance traits are obtained from Q-TARO database [52] and mapped on to the co-expressed modules. We identified 249 QTL genes in our co-expression network that are reported to be associated with resistance/tolerance traits in Q-TARO database. Of these, 73 QTL genes are up-regulated, 67 down-regulated and 109 are not differentially expressed. As shown in **Figure 2**, twenty seven of the 73 up-regulated genes are associated with drought tolerance and mapped to the up-regulated modules, Turquoise (23 genes), Yellow (3) and Tan (1), while thirteen of the 67 down-regulated genes are associated with drought and mapped to the down-regulated modules, Green (7), Blue (4) and Magenta (2). Some of these drought tolerance QTL genes that mapped to the Turquoise module are known transcription factors, *viz*., OsHOX22, bZIP TFs (e.g., OsABF1, OsbZIP23 and ABI5-Like1(ABL1)), and NAC TFs (e.g., OsNAC6 and OsNAC2). Apart from these QTL genes, Turquoise module also harbors QTL genes associated with salinity (21), cold stress (7), other soil stress tolerance (8), blast resistance (8), bacterial blight (6), etc. This indicates common stress responsive mechanisms in abiotic and biotic stress conditions. Previous studies suggest that drought and salinity responses share common pathway components and regulators in the plant [53]. This seems to be well captured in our study with bZIP TFs involved in ABA-mediated stress response under drought stress, kinases (SnRK2-type SAPK4) with a role in ABA signaling and also implicated in the regulation of ion homeostasis and salinity tolerance (Diédhiou et al., 2008), PLDα involved in ABA-regulated stomatal closure, zinc-finger proteins (e.g., ZOS3-21 - C2H) involved in salt tolerance and ABA induced anti-oxidant defense, being up-regulated and co-expressed together in the same module. We also observe some of the QTL genes associated with biotic stress to be up-regulated and co-expressed with drought-responsive genes in Turquoise module. For example, selenium-binding gene OsSBP (associated with increased H_2_O_2_), OsGH3.2 involved in modulating auxin and ABA as well as in pathogen defense by suppressing pathogen-induced IAA accumulation, OsWRKY31 (involved in auxin signaling) and OsNAC6 TFs (involved in broad spectrum of resistance including wounding and blast disease). Similarly, majority of the down-regulated genes associated with blast resistance such as the well characterized WRKY45 and OsNPRI involved in salicylic acid (SA) signaling pathway and defense response, some drought-associated QTLs, e.g., OsGSK1 associated with brassinosteroid (BR)-signaling and flowering, three AP2-EREBP TFs (OsDREB1E, OsDREB6, OsDREB1B) involved in ethylene singling are down-regulated and part of the Green module.

**Figure 2:**
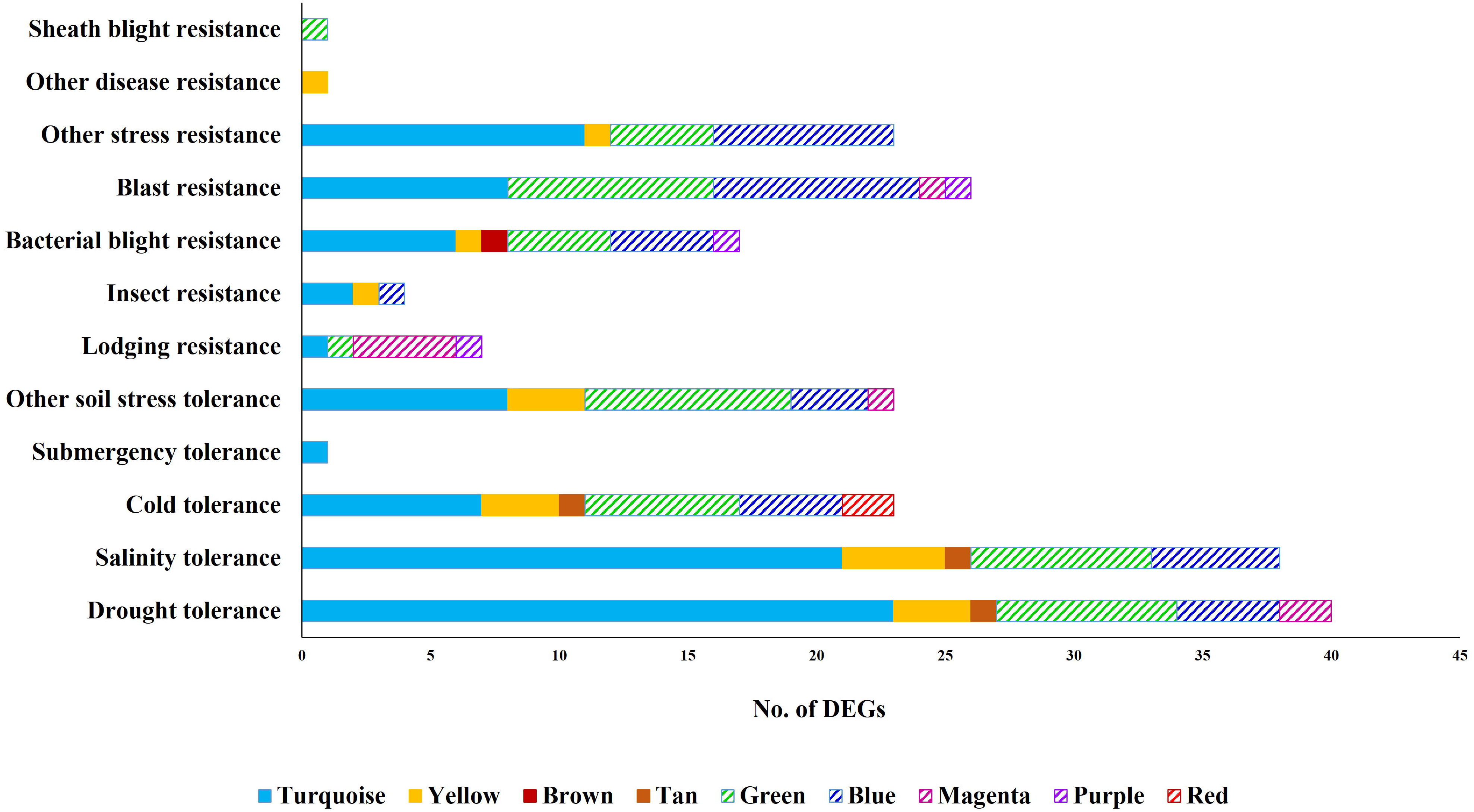
Number of differentially expressed genes (DEGs) from the co-expressed modules that span the QTLs associated with various stress conditions is depicted. The solid bars represent modules with up-regulated genes, while striped bars represent modules with down-regulated genes. It may be noted that a significant number of DEGs from Turquoise, Yellow, Green and Blue modules are associated with osmotic stress (drought and salinity tolerance). The gene ID-QTL mapping is taken from the Q-TARO database.

### Analysis of Minimal Drought-adaptive Modules

To understand the core drought-responsive signaling and metabolic changes, we discuss below the analysis of two PPI networks, uDTN (comprising 466 genes and 1015 interactions) and dDTN (comprising 665 nodes and 8719 interactions). These networks integrate protein-protein interactions with co-expressed gene profiles for the DEGs that are topologically significant (top 20% k_IM_ and k_ME_ values) in the co-expression network. As discussed in Section 1.4, protein-protein interactions for these gene sets are extracted from STRING database and co-function networks, RiceNet and AraNet (**Table S4 and Table S5, Additional File 1**)

### Analysis of uDTN

To identify the processes that are induced in response to drought stress, we applied MCL algorithm to identify tightly coupled gene clusters of genes (size ≥ 5) that are functionally enriched in STRING database for domains, pathways, or GO terms. This resulted in 13 clusters summarized in **Table 4**. A brief description of the biological processes captured by these clusters is given below based on annotations in MapMan [54] and RGAP [55].

**Table 4.** Functional enrichment of gene clusters identified in up-regulated drought tolerant network (uDTN) using MCL algorithm. (TQ: Turquoise, Y: Yellow, BR: Brown, T: Tan).

| Cluster | Associated Biological Processes | Total Genes | Cluster | Associated Biological Processes | Total Genes |
| --- | --- | --- | --- | --- | --- |
| U1 | ABA-signaling and secondary metabolism | 35<br>(23TQ, 10 Y, 2BR) | U8 | Ubiquitination | 8<br>(4TQ, 4Y) |
| U2 | Molecular chaperons and heat stress TFs (Hsfs) | 19<br>(8TQ, 2T, 9Y) | U9 | TCA cycle | 6<br>(5Y, 1BR) |
| U3 | Cell wall and amino acid metabolism | 16<br>(13TQ, 3Y) | U10 | Protein kinases | 5 TQ |
| U4 | Amino acid degradation and mitochondrial ETC | 15<br>(13TQ, 2Y, 1T) | U11 | Mitochondrial ETC/ATP synthesis | 5<br>(3Y, 2TQ) |
| U5 | RNA binding and processing | 13<br>(6BR, 4TQ, 2T, 1Y) | U12 | Starch synthesis and degradation | 5 TQ |
| U6 | Protein degradation | 10<br>(5Y, 5TQ) | U13 | bZIP TFs | 5 TQ |
| U7 | Plant defence system | 9<br>(7TQ, 1Y, 1BR) |  |  |  |

### Cluster U1, U10 and U13: Components of ABA signaling, primary metabolism and biosynthesis of secondary metabolism

From our analysis, we observe that the three clusters, U1, U10, and U13, capture the abscisic acid (ABA) signal transduction process, and components of primary and secondary metabolism. It is known that on perceiving abiotic stress, the adaptive response of plants is controlled mainly by the phytohormone, ABA. Under drought stress, a sudden increase in ABA synthesis is induced accompanied by various physiological and morphological changes in the plant, namely, regulation of growth, stomatal closure, hydraulic conductivity, etc. However, mechanisms of fine-tuning the ABA-levels are not yet well understood. Earlier studies suggest that stress responsive genes mediate through at least two pathways: ABA-dependent and ABA-independent, with a possible crosstalk between their components. Calcium, which serves as a second messenger for various stresses, is proposed to be a strong candidate, mediating such cross talks. Recent progress in our understanding of ABA signal transduction indicates that the central signaling module comprises three protein classes: Pyracbactin Resistance/Pyracbactin resistance-like/Regulatory Component of ABA Receptor (PYR/PYL/RCARs) proposed to be the ABA receptors, Protein Phosphatase 2Cs (PP2Cs) which act as negative regulators, and SNF1-related protein kinases 2 (SnRKs) which are positive regulators. In the presence of ABA, the PYR/PYL/RCAR-PP2C complex formation leads to inhibition of PP2C activity, allowing activation of SnRKs which target membrane proteins, ion channels and transcription factors, and facilitate transcription of ABA-responsive genes (see **Figure 3(A)**, reproduced from KEGG [56].

**Figure 3:**
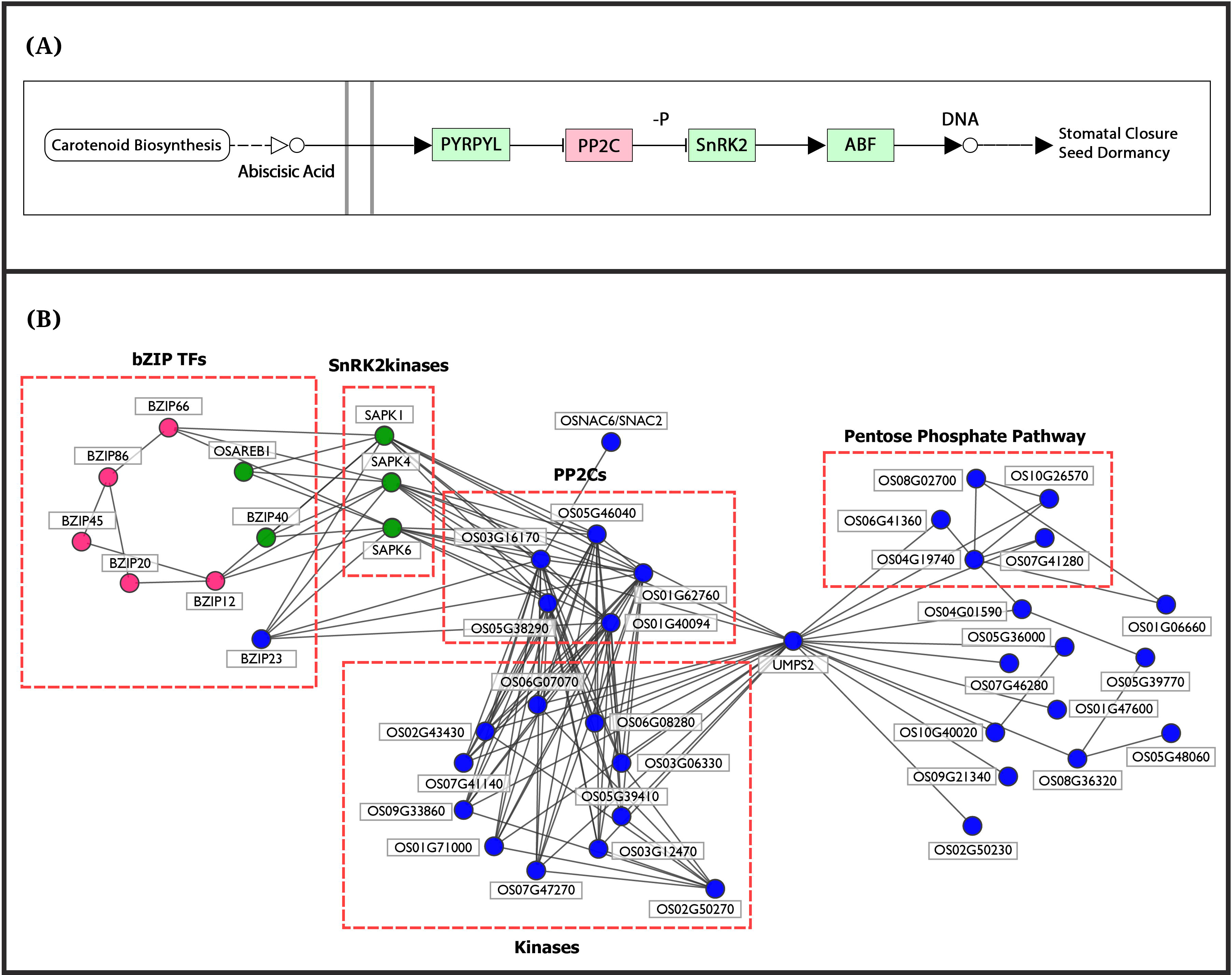
(A) Components of ABA signalosome complex: PYR/PYL receptors, PP2Cs, SnRK2 kinases and ABF/bZIP transcription factors are depicted (reproduced from KEGG). (B) Subnetwork of Cluster U1 (‘blue’), Cluster U10 (‘green’) and Cluster U13 (‘pink’) obtained using MCL algorithm on up-regulated drought-tolerant network (uDTN) is seen to capture the crosstalk between the signaling components: PP2Cs, kinases, bZIP23, OsNAC6, SnRK2 kinases (Cluster U1), bZIPs (Cluster U1 and Cluster U13) with metabolic components of Cluster U1 via UMPS2. This clearly indicates the role of UMPS2 as a novel candidate for drought.

We observe that the subnetwork comprising three gene clusters in **Figure 3(B)**: U1 (35 genes in blue), U10 (5 genes in green) and U13 (5 genes in pink) capture very well essential components of the ABA-signaling pathway (**Figure 3(A)**). Based on functional analysis, U1-cluster genes can be divided into two groups, one involved in signaling processes and the other involved in primary metabolism and biosynthesis of secondary metabolites. The genes involved in signaling processes include 10 kinases, 5 PP2Cs and a membrane-bound ankyrin-like protein (LOC_Os02g50270) with a probable role in calcium-mediated signaling. From AraNet we find that the Arabidopsis ortholog of LOC_Os02g50270 is co-expressed with calmodulin genes, CAM3, 6 and 7. These Ca^2+^ sensors have been implicated in various developmental and environmental stimuli response [57]. The upstream promoter region of the ankyrin-like protein consists of ABRE elements, heat stress-responsive elements, and light responsive elements (e.g., G-box) indicating its possible role in stress. In our subnetwork we observe that its first neighbors are kinases involved in signaling and protein posttranslational modifications. The 10 kinases include cell-wall associated kinase 25, OsWAK25, involved in pathogenic resistance [58], chloroplast localized kinases (LOC_Os07g47270, LOC_Os01g71000 and LOC_Os03g06330) involved in redox regulation and phosphorylation [59], OsRPK1-receptor like kinase, a negative regulator of auxin transport [60] and receptor-like cytoplasmic kinase, OsRLCK-XV (LOC_Os06g07070) highly induced during drought and cold stress [61]. Topologically significant (high degree), the biologically relevant (co-clustered with ABA signalosome and connected to important kinases) along with ABA-responsive *cis*-elements in its promoter and potential role in calcium mediated signaling makes the ankyrin-like protein (LOC_Os02g50270) a likely novel candidate for drought stress. Cluster U1 also consists of two transcription factors (TFs), ABF-type OsbZIP23 and OsNAC6/SNAC2. The OsbZIP23 TF is associated with ABA signaling pathway and strongly induced by drought, high-salinity, polyethylene glycol (osmotic stress) and ABA-treatments. The interactions of OsbZIP23 with 7 genes of U10-cluster and 4 genes of U13-cluster in this subnetwork are in accordance with a genome-wide study using a ChIP assay by Zong et al [62]. Transgenic rice plants overexpressing OsNAC6/SNAC2 displayed increased sensitivity to ABA and tolerance to dehydration and high salinity stresses, suggesting its role as a transcriptional activator in both salinity and drought stresses [63–65] (Hu et al., 2008; Nakashima et al., 2007; Sun et al., 2015). Thus, we see that this subnetwork is able to capture apart from the ABRE regulon, role of NAC regulon in the transcriptional networks of abiotic stress response.

The rice SnRK2 kinases are shown to be over-expressed under abiotic stress with a probable role in phosphorylating ABF family of bZIP TFs for down-stream ABA signaling [67,68]. This association is well captured in our subnetwork exhibiting interactions of OsbZIP23 TF with SnRK2 kinases and PP2Cs (Cluster U1). The 5 ABF genes of Cluster U13 (red) are also seen to exhibit direct/indirect association with PP2Cs (**Figure 3 (B)**) and their role in stress response is well documented. The OsbZIP66/TRAB1 gene encoding ABF-type TF (ABRE-binding bZIP factor) is reported to be up-regulated under dehydration and salt stress conditions [69]. The terminal or nearly terminal event of the primary ABA signal transduction pathway is phosphorylation of this gene [70,71]. The TFs OsbZIP45 and OsbZIP12 are also well characterized for their role in providing drought tolerance (Nijhawan et al, 2008, Joo et al., 2014), while OsbZIP20/RISBZ3/RITA-1 is shown to be expressed in the late stages of seed development with a possible role in regulating seed-specific genes [74,75]. As both seed maturation and response to drought is co-regulated by ABA, the involvement of OsbZIP20 in drought tolerance can be further investigated. Similarly, OsbZIP86 being up-regulated and topologically significant in the co-expression network and its association with other stress-responsive bZIP TFs in the uDTN, suggest its possible role in stress response. These interactions suggest OsbZIP20 and OsbZIP86 transcription factors as probable novel candidates for drought stress.

The 10 kinase signaling genes (Cluster U1) are connected to 16 genes associated with primary metabolism and biosynthesis of secondary metabolites via Uridine 5’-monophosphate synthase, UMPS2 (LOC_Os01g72250). It is a precursor for pyrimidine nucleotides and is associated with the last step of de novo pathway of pyrimidine biosynthesis. Eleven of these 16 genes are involved in the biosynthesis of secondary metabolites associated with sugars and amino acids, and 5 genes belong to Pentose Phosphate Pathway (PPP). Role of PPP is to maintain redox potential to protect against oxidative stress by producing reductants such as NADPH [76]. To ensure these genes are possible targets of ABA-induced downstream signaling cascade, we carried out promoter analysis of these 17 genes (16 metabolic pathway genes+UMPS2). For this, 1 kb upstream region of these genes are analyzed using *cis*-acting regulatory elements database, PlantPAN (v2.0) [77]. The results indicate the presence of ABRE motifs like ‘ABRELATERD1’ in all the 17 genes upstream of the genes (**Table S6, Additional File 1)**. The gene UMPS2 is up-regulated, has ABRE motifs in its promoter region and is topologically significant (‘hub’ gene) both in the co-expressed module and in uDTN. These results suggest the probable role of UMPS2 in relaying ABA-induced metabolic changes, hitherto unknown. To our knowledge there is only one proteomic study in which UMPS2 has been reported to be up-regulated in rice seedlings after ABA treatment followed by subsequent salt stress [78]. Thus, this analysis suggests UMPS2 as a potential drought-tolerant candidate gene in rice.

### Cluster U2: Molecular chaperons and heat stress transcription factors

Almost all stresses induce the production of a group of proteins called heat-shock proteins (Hsps) or stress-induced proteins as adverse environmental conditions disrupt protein folding and lead to an increase in reactive oxygen species (ROS) and oxidative stress within the cell [79,80]. Their transcription is controlled by regulatory proteins called heat stress transcription factors (Hsfs) by binding to the highly conserved heat shock elements (HSEs) in the promoter regions. The Hsps function as molecular chaperones, regulating cellular homeostasis and preventing protein misfolding and aggregation, thus promoting survival under stressful conditions. Cluster U2 in uDTN clearly captures these interactions. It contains 19 genes of which 7 are heat shock proteins (5 DnaJ/Hsp40 family and 2 DnaK/HSP70 family), 4 HsFs (OsHsfA9, OsHsfA2e, OsHsfB2c and OsHsfA2c), MYB TF (LOC_Os04g30890), CDK5RAP3 and another molecular chaperon, ERD1 protein (chloroplast precursor, LOC_Os02g32520) and involved in heat stress. The Hsp70s, assisted by co-chaperone DnaJ-proteins and nucleotide exchange factor constitute a chaperone machine that participates in protein folding, prevention of protein aggregation, translocation of proteins across membranes, targeting proteins towards degradation, and regulation of translation initiation [81,82].

Promoter analysis of the four HSFs revealed presence of ABRE elements suggesting their role in ABA-dependent signal transduction. It has been suggested that Hsps and Hsfs might be important elements in crosstalk of different stress signal transduction networks [83]. The first-degree neighbors of these TFs (extracted from STRING database) mapped to 13 genes in our co-expression network, with 10 genes being common neighbors to all the 4 TFs. Promoter analysis of the up-regulated neighbors (8) was carried out to identify possible regulatory elements in these transcriptionally active genes under drought. Majority of these genes are Hsps (LOC_Os09g29840, two Hsp70s, Hsp90, Hsp40, Hsp81-3, dnak - LOC_Os03g11910) and an RNA recognition motif family protein, LOC_Os03g15890. All the 8 genes have ABRE elements in their promoters, 5 of the genes have heat shock elements (HSEs) and 6 of them also have *cis*-elements which are responsive to methyl jasmonate (MeJa). These results indicate that the Hsps and HSFs are induced in an ABA-dependent manner and at the same time, they are also responsive to jasmonates indicating complex interplay between the two phytohormones. In a study by Singh et al., (2012), it has been shown that OsHsfA2c (part of Turquoise module) binds to OsClpB-cyt/Hsp100 promoter (also part of Turquoise module) and possibly play the role of transcriptional activator in heat stress. Our analysis show that OsHsfA2c exhibits highest fold change, followed by OsHsfB2c, OsHsfA2e and OsHsfA9. Also, both ABRE and HSE elements are present in the promoter of OsClpB/Hsp100 gene, and is up-regulated in our network. These results suggest OsHsfA2c may play the role of transcriptional activator even in drought stress and the HSFs, OsHsfA2c, OsHsfB2c and OsHsfA2e are probable biomarkers for drought tolerance.

### Cluster U3, U4, U9 and U11: Components of Amino Acid Metabolism, Cell Wall and TCA/Citric Acid Cycle and Mitochondrial ETC/ATP synthesis

Maintaining cellular elasticity and integrity by increased cellulose and hemicellulose synthesis is an important drought-tolerant trait [85,86]. We observe that these processes are captured by the subnetwork formed by Clusters U3 (Cell wall and amino acid metabolism), U4 (Amino acid degradation and mitochondrial ETC), U9 (TCA cycle) and U11 (Mitochondrial ETC/ATP synthesis), shown in **Figure 4**.

**Figure 4:**
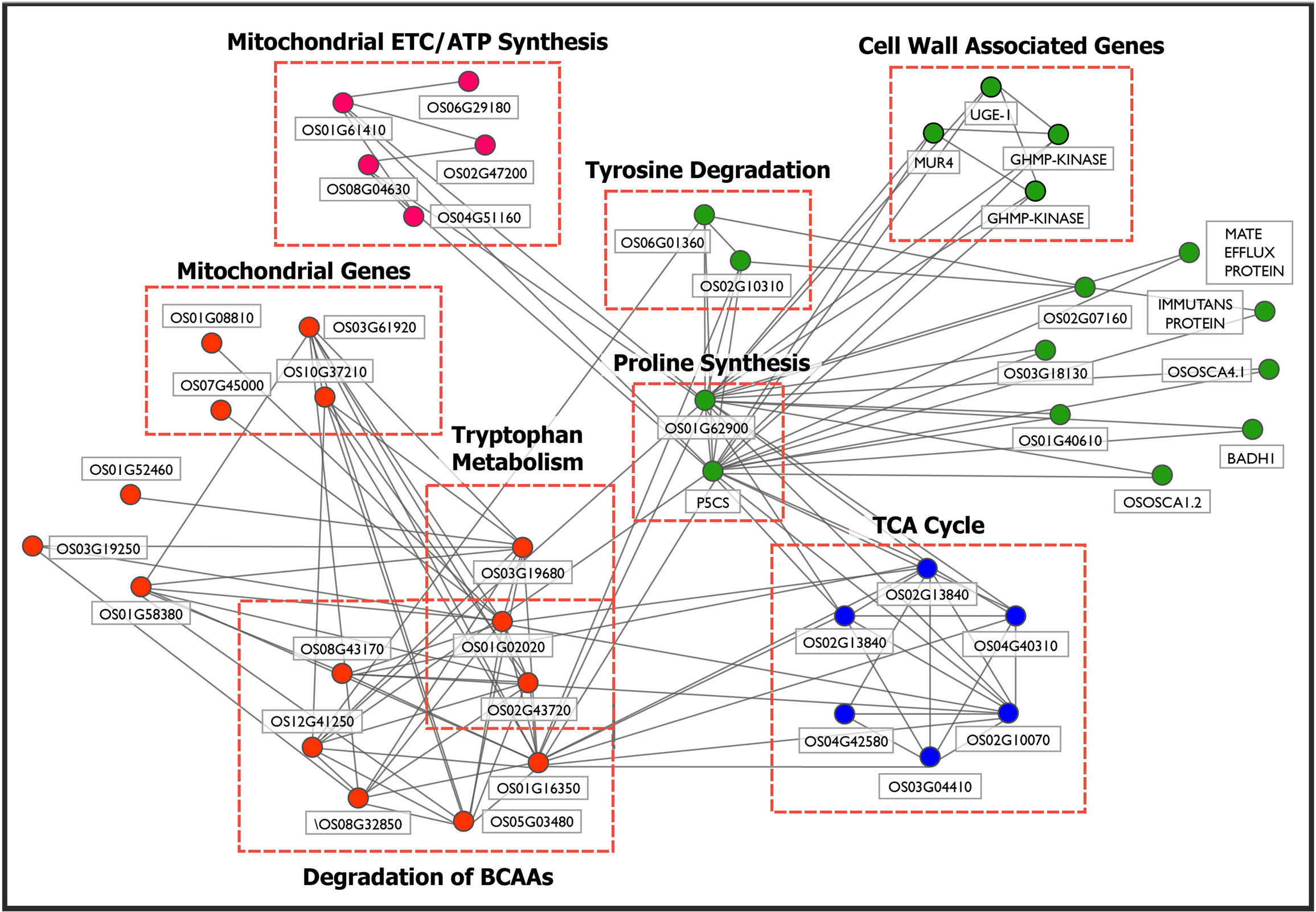
Subnetwork of Cluster U3 (‘green’), Cluster U4 (‘orange’), Cluster U9 (‘blue’) and Cluster U11 (‘pink’) obtained using MCL algorithm on up-regulated drought-tolerant network (uDTN) is shown. It depicts the crosstalk between the components of Amino Acid Metabolism (Cluster U3 and U4), Cell Wall metabolism (Cluster U3), TCA/Citric Acid Cycle (Cluster U9) and ATP synthesis (Cluster U4). These interactions indicate link between structural integrity of plant and cellular energy generating processes as a drought-adaptive mechanism.

It is known from literature that Proline accumulation occurs under various stress conditions and imparts beneficial role to plants by maintaining osmotic balance, stabilizing membranes (thereby preventing electrolyte leakage) and in scavenging of free radical and ROS species [87,88]. Proline is also known to be a key determinant of many cell wall proteins that play an important role in cell wall signal transduction cascades, plant growth and differentiation. The interactions of amino acid kinases P5CS (LOC_Os05g38150) and LOC_Os01g62900, involved in proline biosynthesis with 4 genes involved in cell wall synthesis, OsUGE1 (UDP-glucose 4-epimerase 1), MUR4 (UDP-arabinose 4-epimerase), and GHMP kinases (LOC_Os02g04840 and LOC_Os03g61710) in Cluster U3, capture the role of proline as an important component of the cell wall matrix, providing tensile strength to the cell walls [89]. In **Figure 4**, we observe that the genes involved in proline biosynthesis in Cluster U3 is involved in crosstalks with clusters U4 (Amino acid degradation and mitochondrial ETC), U9 (TCA cycle) and U11 (Mitochondrial ETC/ATP synthesis).

Three genes of Cluster U3, glyoxalase family protein (LOC_Os02g07160), Fumarylacetoacetate (LOC_Os02g10310) and Homogentisate 1, 2-dioxygenase (LOC_Os06g01360) are involved in phenylalanine and tyrosine degradation. Fumarate (also a metabolite of citric acid cycle) and acetoacetate (3-ketobutyroate) are liberated in tyrosine catabolic process. Acetoacetate, on activation with succinyl-CoA, is converted into acetyl-CoA (LOC_Os01g02020 of Cluster U4), which in turn can be oxidized by citric acid cycle or used for fatty acid synthesis. This provides a crosstalk between genes of Clusters U3 (Cell wall and amino acid metabolism) and U4 (Amino acid degradation and mitochondrial ETC). Secondary metabolites such as lignins, flavonoids, isoflavonoids, etc. are usually derived from the catabolic pathway of tyrosine [90]. Accumulation of these phenolic compounds during drought is required for ROS scavenging. Moreover, increased lignin deposition, cell wall stiffness and reduced cell expansion is reported to inhibit plant growth (Alvarez et al., 1997; Mouradov & Spangenberg, 2014; Yoshimura et al., 2007). Several stress-responsive genes such as hyperosmolality-gated calcium-permeable channels which act as osmosensors, OsOSCA1.2 (LOC_Os05g51630) and OsOSCA4.1 (LOC_Os03g04450) (Li et al., 2015), MATE efflux family protein (LOC_Os03g42830) involved in transport (Tiwari et al., 2014), immutans protein (LOC_Os04g57320) involved in electron transport, and asparagine synthetase (LOC_Os03g18130) involved in nitrogen metabolism (Gaufichon et al., 2010) are part of this subnetwork (Cluster U3) and connected to the amino acid kinases P5CS (LOC_Os05g38150) and LOC_Os01g62900, involved in proline biosynthesis suggesting transport and distribution of small metabolites.

The degradation of branched-chain amino acids (BCAAs) is elevated during stress to provide intermediates for TCA cycle and electron donors of the mitochondrial electron transport chain to generate cellular energy [97–99]. This process is captured by the six genes of Cluster U4, involved in the degradation of amino acids valine, leucine and isoleucine. The cluster also includes mitochondrial and peroxisomal genes, explaining the degradation of BCAAs occurring predominantly in these two cellular organelles [100]. Cluster U4 includes genes involved in electron carrier activity, *viz*., EFTA (LOC_Os03g61920), electron transfer flavoprotein subunit alpha-mitochondrial precursor and LOC_Os10g37210, FAD-dependent oxidoreductase domain containing protein (involved in electron transfer to mitochondrial respiratory chain). Membrane-bound cytochrome P450 genes (LOC_Os07g45000 and LOC_Os01g08810) with oxidoreductase activity are up-regulated and part of this cluster. Four out of 5 genes of Cluster U11 are also involved in mitochondrial electron transport chain (ETC)/ATP synthesis. This cluster shows an interaction between rice alternative oxidases OsAOX1b (LOC_Os04g51160), OsAOX1c (LOC_Os02g47200), NADH-ubiquinone oxidoreductases (LOC_Os08g04630, LOC_Os01g61410) and putative erythronate-4-phosphate dehydrogenase domain containing protein (LOC_Os06g29180). A number of roles have been associated with AOXs such as optimization of respiratory mechanisms, replenishment of TCA cycle intermediates [101], preventing excess generation of ROS [102] and shown to be up-regulated in response to biotic and abiotic stress conditions [103–106]. The genes of NADH-ubiquinone oxidoreductase complex in mitochondria are entry point to the respiratory chain from TCA cycle and are involved in ATP synthesis. The up-regulation of ATP synthesis is probably to meet energy demands of the plant to sustain metabolism under drought stress [107]. The putative homolog of LOC_Os06g29180 is an NAD+-dependent formate dehydrogenase in Arabidopsis with a probable role in NADH synthesis [108]. Here too, we observe crosstalk between genes of Clusters U3 and U11 via amino acid kinases P5CS (LOC_Os05g38150) and LOC_Os01g62900 and NADH-ubiquinone oxidoreductases (LOC_Os08g04630, LOC_Os01g61410) linking proline catabolism inside mitochondria with a probable role in ROS production [109]

The TCA cycle takes place in mitochondria in a series of reactions to generate ATP. Five genes of Cluster U9 *viz*., citrate synthases (LOC_Os02g13840, LOC_Os02g10070), dehydrogenases (LOC_Os01g16900, LOC_Os04g40310) and aconitase (LOC_Os03g04410), are involved in the Citrate Cycle. Increase in the expression levels of these genes suggest an increase in ATP production as well as increase in reductants and substrates for non-essential amino acid synthesis like glutamine, proline, arginine, etc. [110–112]. Indeed, we do capture these interactions in the crosstalk between U4 (Amino acid degradation and mitochondrial ETC) and Cluster U9. The interactions between HMG-CoA (LOC_Os01g16350) and genes of the melavonate pathway (acetyl-CoA acetyltransferase (LOC_Os01g02020) and HMG-CoA synthase (LOC_Os08g43170)) in Cluster U4 with the genes of the TCA cycle (Cluster U9) provides a snapshot of the different metabolic fates of the acetyle-CoA, a central metabolite which can be incorporated into the TCA cycle for ATP generation in mitochondria, synthesis of amino acid carbon skeletons or metabolized in the cytosol to produce a plethora of phytochemicals required for growth, development and tolerance towards environmental stress [113,114].

### Smaller uDTN Clusters

Apart from the major processes discussed above, several other smaller clusters identified in uDTN revealed various processes up-regulated in drought-stress, and are summarized in **Table 4**. For example, Cluster U5 is involved in RNA binding and harboring genes with RNA recognition motifs. RNA-processing via post-transcriptional modifications and alternative splicing are important components responsible for fine-tuning the regulatory mechanisms in response to environment stress [115–117]. Clusters U6 and U8 are involved in protein ubiquitination and degradation which are major posttranslational modifications, providing a dynamic and reversible control over processes such as degradation of misfolded proteins, trafficking, signal transduction and cell division [118,119]. Cluster U7 consists of genes associated with plant defense system. Other than stress-induced lipases and transporters, genes like CXE carboxylesterase (LOC_Os06g11090) and GID1L2-gibberellin receptors (LOC_Os11g13670 and LOC_Os05g33730) which are involved in biodegradation of xenobiotics (herbicides) are also present in this cluster. These genes are induced as a result of increased oxidative stress in the system [54] and may have a role in ROS scavenging in the plant. Starch synthesis and degradation is an important aspect regarding metabolic shifts during drought stress as seen in Cluster U13. It harbors alpha and beta-amylases involved in breakdown of starch and produce soluble sugars, OsDPE1 involved in transitory starch breakdown and AGPlar involved in starch synthesis. The increase in amylase activity in the shoot can be seen as an important drought tolerant trait, providing tensile strength and maintaining cellular membranes [120,121]. Thus we show that by using an integrated approach by combining co-expression and protein-protein interaction information we are able to capture some of the core biological processes in drought response. In the up-regulated PPIN, we are able to extract some of the key components of the ABA signalosome including the PP2Cs, the SnRK2 kinases and a number of ABF-type bZIP transcription factors. Along with the genes involved in signaling, we observe a crosstalk with the genes involved in metabolic pathways like Pentose Phosphate Pathway and biosynthesis of sugars and amino acids via the gene UMPS2, a novel candidate for drought. We also extracted a cluster comprising molecular chaperons and heat shock transcription factors as well as an MYB transcription factor co-clustered with these genes. The other major crosstalk is observed between processes, namely biosynthesis of proline which is an osmolyte and an important component of cell wall matrix, cell wall metabolism, degradation of branched chain amino acids to serve as intermediates for TCA cycle, enzymes of the TCA and finally, mitochondrial electron transport/ATP synthesis to generate cellular energy during stress. There is a re-partitioning of metabolites during drought stress as genes involved in starch biosynthesis as well as degradation are seen to be up-regulated. Finer regulatory mechanisms such as RNA-processing via post-transcriptional modifications and alternative splicing are captured in this network. Finally, we observe that protein ubiquitination and degradation is a prominent process in drought by which degradation of misfolded proteins due to stress, trafficking, signal transduction and cell division are dynamically controlled during stress.

### Analysis of dDTN

A similar analysis is carried out on the down-regulated drought tolerant network, dDTN, to identify the processes down-regulated in response to drought stress by the proposed network-based approach. Seventeen tightly-coupled gene clusters are identified using MCL algorithm in dDTN. Functional annotation for biological processes associated with these gene clusters using MapMan and RGAP is summarized in **Table 5** and is discussed below in detail.

**Table 5.** Functional enrichment of 17 gene clusters identified in down-regulated drought tolerant network (dDTN) using MCL algorithm. (B: Blue, P: Purple, G: Green, M: Magenta, S: Salmon).

| Cluster No. | Function | Total Genes | Cluster No. | Function | Total Genes |
| --- | --- | --- | --- | --- | --- |
| **D1** | Photosynthesis and associated process | 262<br>(178B, 72R, 5S, 3P, 4G) | **D8** | alpha-Linolenic acid metabolism | 7<br>(4B, 3G) |
| **D2** | Cell Wall metabolism | 45<br>(43M, 1B, 1G) | **D9** | Glutathione S-transferase | 6<br>(4G, 2B) |
| **D3** | Signaling and post-translational modifications | 37<br>(22G, 10B, 2R, 1M, 1P, 1S) | **D10** | Serine Metabolism | 6<br>(4B, 1P, 1M) |
| **D4** | Ribosome biogenesis | 25<br>(12B, 8R, 4P, 1S) | **D11** | DNA-repair processes | 5<br>(3B, 2P) |
| **D5** | tRNA biogenesis | 22<br>(9B, 4G, 3R, 1S, 5P) | **D12** | Vitamin B6 metabolism | 5<br>(2B, 2G, 1R) |
| **D6** | Development and Flowering | 11<br>(4B, 7G) | **D13** | Cell vesicle transport | 5<br>(2G, 2M, 1B) |
| **D7** | Signaling and Transport | 8<br>(5B, 1M, 1S, 1G) | **D14** | Biotic Stress | 5<br>(3G, 1B, 1P) |

### Cluster D1: Photosynthesis and associated processes

Photosynthesis, a part of the primary metabolic process in plant, plays a central role in drought response. With decrease in carbon uptake due to stomatal closure, major metabolic shifts take place to re-partition the photo-assimilates between root and shoot, which lead to the cessation of shoot growth and maintenance of root growth under water deficit conditions [122]. Around ∼23% of Cluster D1 genes are directly involved in photosynthesis, *viz*., genes associated with photosynthetic light reaction such as ATP synthases and NADPH dehydrogenase subunits, chloroplast precursors and chlorophyll A-B binding proteins. Closely associated processes with photosynthesis such as carbon fixation, tetrapyrrole synthesis and carbohydrate metabolism are also components of this cluster. Closely clustered with the photosynthetic genes, around 30% genes are involved in protein synthesis, folding, targeting, posttranslational modifications and degradation. On closer look, a large majority of them are ribosomal proteins, many of which are located in chloroplast (21 genes). Genes associated with RNA processing (helicases), regulation and binding (52 genes) are also seen to be clustered together with photosynthetic processes. The crosstalk between these processes suggests down-regulation of photosynthesis and transcription and translation of plastidial genes.

Twenty one genes in this cluster have no functional annotation. These genes are queried in AraNet and the top functional predictions for each gene are listed in **Table S6, Additional File 1.** Of these, ∼20 genes have predicted functions associated with photosynthetic machinery, pathogen response or both. Promoter analysis of these 21 uncharacterized genes using PlantPan database revealed presence of cis-elements like ‘GT1CONSENSUS’ and ‘CGACGOSAMY3’ indicating the presence of light-responsive elements like G-box in all the genes. These elements are known to regulate light-specific gene expressions and response to stimuli [123]. Other motifs like “SITEIOSPCNA” which is shown to be involved in gibberellin-responsive pathway in regulating photosynthesis [124] are also present. The above analyses indicate that the 21 uncharacterized genes of this cluster that are topologically significant and tightly clustered with photosynthetic genes, are likely to be regulated by light, respond to pathogen invasion/wounding and may be influenced by hormones such as gibberellin. We presume these genes to be important candidates for transgenic studies to elucidate their role in photosynthesis and drought-response.

### Cluster D2, Cell Wall Metabolism

Plant cell walls are of critical importance and provide cell shape and mechanical support to withstand the turgor pressure. However, during water deficit, aerial growth is limited due to reduced cell division in the meristematic zones [125] followed by reduced ability of cells to expand the polysaccharide network. Cluster D2 genes are found to be associated with cell wall metabolism, e.g., cellulose synthases OsCESA6, 7 and 8 and cellulose synthase-like genes OsCLSA1 and OsCLF6, and OsBC1L4/OsCOBRA involved in cellulose biosynthesis. While OsCESA6, 8 and OsBC1L4/OsCOBRA are reported to be involved in primary cell wall biosynthesis [126], OsCESA7 is a secondary cell wall-specific cellulose synthase [127]. The gene OsCLF6 is shown to mediate mixed-linkage glucans (MLG) synthesis in rice [128]. MLG is a cell wall polysaccharide, known to be involved in the regulation of cell wall expansion in young tissues [129,130]. This cluster also includes 3 Fasciclin-like arabinogalactan proteins (OsFLA6, 16 and 24) that are involved in cell wall adhesion, signaling, biosynthesis and remodeling and in the development of new shoot and root meristems [10]. Among the 5 uncharacterized genes, we observe that Arabidopsis orthologs of LOC_Os06g11990 and LOC_Os05g32500 are involved in cellulose synthesis while that of LOC_Os07g43990 has a role in transferring glycosyl groups (**Table S3, Data Sheet 1)**. Since cell walls are also the first line of defense against pathogens, some of the genes, e.g., Harpin-induced protein 1 (LOC_Os02g33550) and CHIT1 (LOC_Os09g32080-Chitinase family protein) are down-regulated and are part of this cluster. Some of the kinases (LOC_Os02g43870, LOC_Os03g12250 and LOC_Os04g55620) have LRR domains and are important for protein-protein interactions, signaling and response to biotic stress. Thus, a low turgor pressure during drought stress leads to reduction of growth by the inhibition of cell wall elongation. The down-regulation of Cluster D2 genes clearly reveals the processes associated with cell wall biosynthesis that are down-regulated in leaves.

### Cluster D3, D7 and D14: Down-regulation of biotic stress associated genes

The natural phenolic compound, salicylic acid (SA) is one of the main hormones involved in endogenous signaling in plant pathogen response and disease resistance [131]. In many studies drought tolerance and disease resistance pathways are reported to be antagonistic and the role of ABA in suppressing SA-signaling pathway [132–134]. A transcriptional activator NPR1 is an important regulator of SA-mediated defense signaling pathway and responsible for activation of a large number of downstream pathogen-responsive genes [135,136]. It is shown that ABA suppresses transcriptional up-regulation of WRKY45 and OsNPR1, the two important components of SA-signaling pathway [137]. Also, OsNPR1 has been implicated with reduced drought tolerance in rice [138]. Both WRKY45 and OsNPR1 (part of Cluster D3) are co-expressed in Green module and down-regulated in agreement with earlier studies, thus capturing the crosstalk between ABA and SA signaling pathways.

Majority of Cluster D3 genes (∼ 23) are kinases including receptor-like kinases involved in signaling and post-translational modifications. Many of these genes are shown to be involved in plant immunity against pathogens and are potential pattern recognition receptors (PRRs), e.g., leucine-rich repeats (LRR) and ankyrin (ANK) repeats [139–142]. This module consists of three LRR repeat genes, Strubbelig-receptor FAMILY8 precursor (LOC_Os06g42800), a putative transmembrane protein kinase 1, a putative protein kinase (LOC_Os01g41870) and Brassinosteroid insensitive 1-associated receptor kinase 1 precursor (BAK1, LOC_Os11g39370). Previous studies indicate that these kinases influence cellular morphogenesis, orientation, cellular proliferation, size and shape [143–145]. Genes with ankyrin repeats include OsNPR1 (LOC_Os01g09800) a master regulator characterized for resistance to plant pathogens [146–148] and LOC_Os03g63480, are both involved in biotic stress and down-regulated in the network.

Three disease resistance proteins *viz*., RGA3, RPM1 and LOC_Os12g25170 with NB-ARC domains containing LRR repeats and having ATPase and nucleotide binding activities are part of Cluster D14. These proteins are involved in pathogen resistance and help in discriminating between self and non-self (R-protein mediated response) [149]. Cluster D7 consists of 2 genes (OsGF14b and OsGF14e) that belong to 14-3-3 gene family. These signaling proteins are reported to be involved in biotic stress and probably associated with JA and SA pathways [150,151]. They interact with Beta-2-and Beta-5-tubulin proteins which play critical role in cell division, elongation and growth. Binding of 14-3-3-signaling proteins with those involved in cellular organization (tubulins) hint at finer regulations of cell cycle events [152]. Thus, the analysis of the subnetwork of Cluster D3, D7 and D14 indicate suppression of the defense system of plants under drought stress, that is, the vulnerability of the plant to pathogen infection under drought condition.

### Cluster D4 and D5: Biogenesis of rRNA and tRNA genes

A number of clusters are associated with synthesis, post-translational modifications and processing of ribosomal proteins. Cluster D4 includes genes involved in pre-rRNA processing in nucleolus and cytoplasm (nucleolar protein 5A-LOC_Os03g22880, LOC_Os04g49580 and Gar2-LOC_Os08g09350), ribosomal proteins (LOC_Os08g41300, LOC_Os06g03790, LOC_Os06g02510 and LOC_Os03g41612), DEAD/DEAH box helicases and WD domain containing proteins, t-RNA synthetases (LOC_Os05g08990 and LOC_Os04g02730) and other RNA-binding genes. The rRNA processing and ribosome biogenesis is tightly coupled with growth and cell proliferation. Down-regulation of these pathways hint at conservation of energy by the plant [153,154]. Closely interacting with Cluster D4, is Cluster D5 with genes involved in amino-acyl tRNA biosynthesis (8 genes), genes involved in protein-processing like ubiquitin conjugating enzymes (LOC_Os01g46926, LOC_Os04g58800, LOC_Os06g44080 and LOC_Os03g03130), further indicating a decrease of protein turnover, cell proliferation and growth.

### Cluster D6: Development and Flowering

Cluster D6 consists of zinc finger domain containing proteins-LOC_Os04g46020, OsTIFY1b, OsBBX25, zinc finger protein-LOC_Os08g15050, MADs box genes (OsMADS50 and OsMADS55) and response-regulator genes (OsRR10 type-A and OsRR2 type-A). The TIFY family genes are involved in development and have been shown to be stress-responsive [155]. The MADs box and BBX genes are also involved in flowering and development [156–158]. Interestingly, rice response regulator (OsRR) genes are part of the cytokinin-dependent signal-transduction pathways which also affect development and morphological changes in the plant [159,160]. The down-regulation of genes in this cluster suggests a decrease in growth, development and delay in flowering of the plant under drought stress.

### Cluster D8: α-Linolenic acid metabolism

The phytohormone Jasmonic Acid (JA) regulates a wide array of biological processes in plants including growth, development, secondary metabolism, pathogen defense, wounding as well as tolerance to abiotic stress [161]. Two genes of GH3 family, *viz*. OsJAR1 and OsJAR2 control the homeostasis of JA by forming bioactive JA-amino acid conjugates. They have been reported to be involved in wound and pathogen-induced JA-signaling [162]. Also, OsOPR7, LOX7 and OsAOS1 genes that are involved in JA synthesis [163,164] are down-regulated and part of this cluster. The wound inducible and leaf-specific gene, OsHPL3 of this cluster is reported to modulate the amount of JA by consuming common substrates [165]. As majority of the genes in JA biosynthesis are down-regulated, it can be inferred that during drought stress, there is an increased susceptibility to wounding and pathogen attack.

### Other smaller dDTN Clusters

Apart from some of the major clusters discussed above, several smaller clusters are detected (**Table 5**). For example, Cluster D9 consists of six glutathione s-transferases (GSTs). The GST family of genes are widely involved in cellular detoxification and can act on substrates such as endobiotic and xenobiotic compounds [166]. However, studies have indicated that not all the GST genes express the same way across different stress, tissue and developmental conditions [167]. A similar behavior is observed in the present case with GSTs in D9 are down-regulated, but 12 other GSTs are up-regulated in the overall co-expression network, and are not part of uDTN. Cluster D10 consists of genes associated with serine metabolism and are down-regulated. Among these, two genes are phosphoserine phosphatase which catalyzes the last step in the plastidial phosphorylated pathway of serine biosynthesis. The other genes include serine hydroxymethyltransferase (SHMT) gene and three aminotransferases with a probable role in glycine cleavage system. The glycine cleavage system is required for photorespiration in C3 plants and closely interacts with serine SHMT for the methylation of glycine to serine. These amino acids have been associated with photorespiration and precursors for other metabolites involved in stress protection [168]. It has been shown that DNA repair mechanisms are affected by drought stress. This is captured by Cluster D11 genes involved in maintaining genome stability and repair mechanisms. With increase in ROS due to stress, plants undergo ABA-mediated growth retardation with increase in DNA damage, inhibition of DNA replication and cell division [169–171]. Genes involved in vitamin B6 metabolism are captured in Cluster D12. Lower levels of vitamin B6 levels have been attributed to impairment of flowering and reduced plant growth in Arabidopsis mutants [172,173]. The Cluster D13 consists of genes 3 major sperm proteins (MSPs) which have not been studied in the context of stress response. These MSPs have a probable role in cell vesicle transport and mediate protein-protein interactions within or between cells [167]

## Discussion

The fascinating complexity of biological processes in plants is reflected in the intricate interactions in co-expression networks. With the availability of a large number of transcriptomic datasets in public repositories, co-expression network analysis allows us to integrate this information and perform large-scale multi-genic studies. On one hand, we need this complexity to infer functional annotations of genes from tightly connected neighborhood (‘guilt-by-association’), while on the other hand, we need to dissect the networks to remove noisy connections and identify critical biological processes. Various approaches have been proposed, e.g., construction of condition-dependent co-expression networks with context-specific interactions [174,175], split co-expression networks into functional modules which may represent tightly coordinated biological processes [176,177], use of functional and comparative genomics [178,179], etc.

The simple differential expression analysis using fold-change and *t*-test, commonly carried out in meta-analytic studies, generally result in a large number of DEGs. For instance, out of 14,270 genes, we observe 6,454 genes (∼ 44%) to be differentially expressed in our study. A detailed functional analysis of such a large number of genes is not feasible, many of which may be false positives. Also, since a typical meta-analytic study based on publicly available microarray datasets is likely to have heterogeneity in the data arising from various sources such as microarray platforms, tissues, developmental stages, experimental conditions, etc. For the analysis, DEGs are restricted to only the 10 drought-responsive modules, which are further filtered based on topological measures, *viz.*, intramodular connectivity (K_IM_) and eigengene-based centrality (K_ME_) to identify biologically significant genes across the genotypes. The DEGs are ranked by K_IM_ and K_ME_ values and top 20% genes from each of the 10 drought-responsive modules are considered, resulting in 1,647 DEGs (754 up-regulated and 893 down-regulated). We observe that greater than 50% of these topologically significant DEGs are well represented in majority of the drought-tolerant genotypes considered (**Table S3, Additional File 2**), clearly suggesting the biological significance of these genes across different stages/genotypes of the plant. This filtered set of DEGs is used in the construction of protein-protein interaction networks, uDTN and dDTN, by extracting known protein-protein interactions (using various resources, *viz*., STRING, AraNet and RiceNet) within the set of up-regulated and down-regulated genes, respectively. Tightly-coupled gene clusters are identified in the two PPI networks, uDTN (466 nodes, 1015 edges) and dDTN (665 nodes, 8719 edges) to capture various stress-responsive processes. It is confirmed that these gene clusters are well-represented across all the 9 data subsets (**Table S5 (a) and S5 (b), Additional File 2**) considered. Thus, by the proposed approach, drought-responsive processes commonly induced/repressed across various drought-tolerant genotypes are identified and analyzed.

Plant’s response to drought stress involves complex interactions across several layers leading to molecular, biochemical, physiological and morphological changes in the plant. By analyzing the PPINs, we show that the analysis of tightly coupled gene clusters enable us to capture many important molecular and biochemical changes occurring in response to drought. The ABA signaling machinery and the interaction of its signaling components with the metabolic pathways is seen at the forefront of the up-regulated processes. We observe cross-talks between clusters involved in ABA-signaling and secondary metabolism (U1), protein kinases (U10) and bZIP TFs (U13) which are up-regulated across all the three stages of plant (**Table S5 (a), Additional File 2**). We also observe cross-talk between genes involved in metabolic pathways e.g., Pentose Phosphate Pathway and biosynthesis of sugars and amino acids via the ‘hub’ gene UMPS2, which is identified as a novel candidate gene for drought tolerance. These interactions observed among the signaling and metabolic components are ubiquitous across all the drought tolerant data subsets considered.

Heat shock proteins (Hsps) function as molecular chaperons and prevent protein misfolding and aggregation. The presence of ABRE *cis-*elements in the promoters of 4 HsFs (OsHsfA9, OsHsfA2e, OsHsfB2c and OsHsfA2c) and their first neighbors (in uDTN) clearly emphasizes that these are induced in an ABA-dependent manner during drought stress. The protective role of (Hsps) in maintaining cellular homeostasis during drought stress is well-captured by Cluster U2 of uDTN. Further, presence of *cis*-elements associated with heat shock elements in promoters of their neighbors and responsive to methyl jasmonate (MeJa) indicate that depending upon the environment, they can function as molecular switches during stress. We observe 8 data subsets have ≥ 50% representation of genes of this cluster (except IRAT109 with 42.1%), and greater than 80% in Dagad deshi-seedlings and DK151-leaves in panicle elongation and booting stages). The gene OsHsfA2e, which is known to enhance tolerance to environmental stresses is up-regulated across all the 9 data subsets [180], while OsHsfA2c with highest fold-change is up-regulated in 6 datasets (**Table S7, Additional File 1**).

The other major crosstalk is observed between processes such as biosynthesis of proline (an important component of cell wall matrix, Cluster U3), cell wall metabolism and degradation of BCAAs (Cluster U4) to serve intermediates for TCA cycle (Cluster U9) and mitochondrial electron transport/ATP synthesis (Cluster U11) corroborating that TCA cycle provides carbon skeletons for the synthesis of amino acids like proline through α-Ketoglutarate/Glutamate-linked aminotransferase reaction [181]. The cluster U3 involved in cell wall metabolism and amino acid metabolism is well represented across all 9 data subsets (62.5% - 100%). Interestingly, both the genes involved in proline biosynthesis (P5CS (LOC_Os05g38150) and LOC_Os01g62900) is up-regulated across all the 9 data subsets highlighting the role of proline as an osmoprotectant across all stages of the plant. The cell wall associated gene OsUGE1 (UDP-glucose 4-epimerase 1) is also up-regulated across all the data subsets to provide metabolites for cell wall biosynthesis. Only one gene of Cluster U9 (TCA cycle) is observed to be up-regulated in Azucena and IRAT109 datasets indicating that these two genotypes may have less efficient methods of ATP generation and substrates for non-essential amino acids compared to other genotypes. Particularly, the genes NAD-dependent isocitrate dehydrogenases (OsIDHa and OsIDHc) are down-regulated in Azucena while citrate synthase (LOC_Os02g10070) and aconitate hydratase (LOC_Os03g04410) are down-regulated in IRAT109.

From the analysis of uDTN clusters in **Table S5 (a), Additional File 2** we observe that genes of Cluster U5 (RNA binding and processing) do not exhibit similar expression pattern across all the 9 data subsets. The 3 data subsets from seedlings stage (23.1% - 46.2%) and DK151 (leaves in panicle elongation stage, 46.2%) exhibit fewer DEGs. The reason for this variation could be this cluster comprises genes having RNA-binding motifs that are involved in diverse processes such as splicing, repair and synthesis of nucleotides, which may vary across various stages of plant. An analysis of 9 data subsets across all the clusters shows that N22 (seedlings) and IRAT109 datasets have overall fewer up-regulated DEGs, 37.3 % and 43% respectively, (**Table S1** and **Table S2, Additional File 2**). This indicates that the RNA-mediated processes captured in this analysis are not well represented across various genotypes.

A similar analysis of the down-regulated PPIN (dDTN) revealed photosynthesis as the most significant down-regulated process, along with reduction in protein synthesis in chloroplasts. Among the 9 data subsets, all the genes of photosynthetic cluster are down-regulated in DK151 (panicle elongation stage) followed by 94.6% genes in Vandana (seedlings) and 86% genes in DK151 (booting phase). This clearly indicates that photosynthesis is highly affected in these stages/genotypes and could be a probable drought escape mechanism. On the other hand, only ∼28% of these genes are down-regulated in Bala genotype, which probably hints at efficient photosynthetic machinery, providing a metabolic advantage to this genotype. Specifically, among the 60 genes directly associated with photosynthesis, only 19 are observed to be down-regulated in Bala. The photosynthetic cluster also contains 21 genes with no functional annotation. Functional characterization and promoter analysis suggests these genes to be light responsive genes, leaf-specific, associated with pathogen response and responsive to environmental stimuli (**Table S6, Additional File 1**).

Genes involved in cellulose synthesis, cell wall metabolism and growth (Cluster D2) are down-regulated as expected under drought stress. Membrane associated genes, kinases having LRR domains and involved in biotic stress signaling are co-clustered indicating shared cellular components. Moreover, two of the five uncharacterized genes in this cluster are found to be associated with cellulose synthesis (LOC_Os06g11990 and LOC_Os05g32500) and LOC_Os07g43990 gene has a probable role in transferring glycosyl groups. Very few genes of Cluster D2 are down-regulated in Azucena (4.4%) and Bala (2.2%). On closer inspection, it is observed that most of these genes are not significantly differentially expressed in Azucena. In Bala genotype, majority of these genes (OsCESA6, 7 and 8, OsCLSA1, OsCLF6 and OsBC1L4/OsCOBRA) and Fasciclin-like arabinogalactan proteins involved in cell wall adhesion and signaling are significantly up-regulated, indicating better mechanical support and continued growth in shoot tissues under drought stress.

Drought followed by ABA signaling indicates the shutdown of NPR1 and WRKY45-dependent SA signaling pathway leading to the down-regulation of biotic stress-responsive genes (Cluster D3, D7, and D14). A number of membrane-bound signaling proteins with LRR and ANK repeats with probable role in cellular morphogenesis, cell division and pathogen response are also found to be down-regulated in these clusters. Majority of the signaling and transport related genes (Cluster D7) are down-regulated across all the 9 data subsets indicating susceptibility to biotic stress as well as growth arrests.

Genes of Cluster D8 that capture the role of JA synthesis are also down-regulated leading to decreased levels of JA, indicating an increased susceptibility to wounding and pathogen attack. The overall analysis of dDTN clusters in **Table S5 (b), Additional File 2** indicate that Cluster D4 (ribosome biogenesis) and Cluster D11 (DNA-repair processes) exhibit poor conservation of differential gene expression across the 9 data subsets. Interestingly, in the genotype DK151 at tillering stage, ribosome biogenesis and DNA repair processes are poorly represented, indicating that protein synthesis might be active in this stage/genotype even under drought. On the other hand, all the genes of these two clusters are down-regulated in DK151-panicle-elongation stage and Dagad deshi-seedlings indicating that protein synthesis and DNA repair processes are most affected in these genotypes.

The integrated network-based approach helped in identifying important processes which are well-represented in majority of the data subsets. Moreover, *in silico* functional annotations of uncharacterized genes, e.g., membrane-bound ankyrin-like protein (LOC_Os02g50270) associated with ABA signaling in uDTN, uncharacterized genes associated with known photosynthetic genes and light-responsive promoters, as well as those associated with cellulose synthesis are identified as important drought-responsive biomarkers. Further analysis of these genes using transgenic studies may help in elucidating their role in drought response.

## Conclusion

In this study, a comprehensive analysis of the complex drought-responsive processes using an integrated, systems-level approach across seven drought tolerant rice genotypes is presented. Preliminary analysis of the co-expression network revealed that transcriptional regulatory processes, post-translational modifications followed by photosynthesis are at the forefront of drought response. With the integration of protein-protein interactions, a detailed view of the system is possible and critical processes which otherwise lay hidden are brought forth. The phytohormone abscisic acid is seen to play a central role in regulating the energy metabolism of cells. Protective mechanisms like the transcriptional activation of molecular chaperons, amino acid metabolism and cellular respiratory processes are enhanced. The down-regulation of photosynthetic machinery and cellulose-associated genes explains a cessation of aerial growth during drought stress. The antagonistic relationship observed between ABA and SA-dependent pathogen response pathways indicate increased susceptibility of plants to certain pathogens during drought.

The systems-based approach discussed above, apart from capturing important stress-responsive processes, also help in *insilico* functional annotation of uncharacterized genes. Based on neighborhood analysis in the co-expression and protein-protein interaction networks, important biomarkers with significant topological properties and connectivity to other stress-responsive genes have been identified. A systematic dataset-specific analysis revealed that certain processes are well-conserved across all genotypes/stages, while some exhibit genotype-specific variations. Drought resistance/tolerance is indeed a complex process with dependencies on many variables, *viz*., soil, weather conditions, duration of stress, tissue-specific response, etc. Better designed experiments with larger sample sizes and taking these variables into consideration will further enhance the efforts in reliable detection of drought-tolerant biomarkers.

## Declarations

### Availability of data and materials

The data sets in this study are available as Supplementary files.

### Authors’ contributions

SS and NP conceived the study, designed the computational method, analyzed the data, and wrote the manuscript. All authors have read and approved the final manuscript.

### Ethics approval and consent to participate

Not Applicable

### Consent for publication

Not Applicable

### Electronic supplementary material

Table S1, Additional File 1: Dataset details which are used in the construction of network

Table S2, Additional File 1: Co-expression network with genes and module labels.

Table S3, Additional File 1: Module preservation statistics

Table S4, Additional File 1: Up-regulated PPIN (uDTN)

Table S5, Additional File 1: Down-regulated PPIN (dDTN)

Table S6, Additional File 1: Analysis of cis-elements for some of the clusters

Table S7, Additional File 1: Stage-specific DEGs across the datasets

Table S1, Additional File 2: Distribution of DEGs in the 6 microarray studies across 9 data subsets

Table S2, Additional File 2: Distribution of DEGs from up and down-regulated drought-responsive modules in the 6 microarray studies across 9 data subsets.

Table S3, Additional File 2: Distribution of the DEGs (after screening for important genes) used for the construction of uDTN and dDTN shown across 9 data subsets.

Table S4, Additional File 2: Distribution of uDTN and dDTN DEGs shown across 9 data subsets (after extracting PPIs).

Table S5 (a), Additional File 2: Distribution of DEGs in uDTN clusters.

Table S5 (b), Additional File 2: Distribution of DEGs in dDTN clusters shown.

## References

1. Current Conditions | Global Drought Information System. [cited 2017 Apr 5]. Available from: https://www.drought.gov/gdm/current-conditions

2. Rhee SY, Mutwil M. Towards revealing the functions of all genes in plants. Trends Plant Sci. 2014 [cited 2017 Apr 2];19:212–21.

3. Jagadish SVK, Muthurajan R, Oane R, Wheeler TR, Heuer S, Bennett J, et al. Physiological and proteomic approaches to address heat tolerance during anthesis in rice (Oryza sativa L.). J Exp Bot. Oxford University Press; 2010 [cited 2017 Apr 2];61:143–56.

4. Lenka SK, Katiyar A, Chinnusamy V, Bansal KC. Comparative analysis of drought-responsive transcriptome in Indica rice genotypes with contrasting drought tolerance. Plant Biotechnol J. Blackwell Publishing Ltd; 2011 [cited 2017 Apr 2];9:315–27.

5. Ghimire KH, Quiatchon LA, Vikram P, Swamy BPM, Dixit S, Ahmed H, et al. Identification and mapping of a QTL (qDTY1.1) with a consistent effect on grain yield under drought. F Crop Res. 2012 [cited 2017 Apr 2];131:88–96.

6. Yue B, Xue W, Xiong L, Yu X, Luo L, Cui K, et al. Genetic Basis of Drought Resistance at Reproductive Stage in Rice: Separation of Drought Tolerance From Drought Avoidance. Genetics. 2006 [cited 2017 Apr 3];172.

7. Mu P, Li Z. Correlation analysis and QTL mapping of osmotic potential in japonica rice under upland and lowland conditions. Can J Plant Sci. Agricultural Institute of Canada; 2013 [cited 2017 Apr 3];93:785–92.

8. Sims AH, Smethurst GJ, Hey Y, Okoniewski MJ, Pepper SD, Howell A, et al. The removal of multiplicative, systematic bias allows integration of breast cancer gene expression datasets - improving meta-analysis and prediction of prognosis. BMC Med Genomics. BioMed Central; 2008 [cited 2017 Apr 3];1:42.

9. Shabalin AA, Tjelmeland H, Fan C, Perou CM, Nobel AB. Merging two gene-expression studies via cross-platform normalization. Bioinformatics. 2008 [cited 2017 Apr 3];24:1154–60.

10. Johnson WE, Li C, Rabinovic A. Adjusting batch effects in microarray expression data using empirical Bayes methods. Biostatistics. Oxford University Press; 2007 [cited 2017 Apr 3];8:118–27.

11. Christie N, Myburg AA, Joubert F, Murray SL, Carstens M, Lin Y-C, et al. Systems genetics reveals a transcriptional network associated with susceptibility in the maize-grey leaf spot pathosystem. Plant J. 2017 [cited 2017 Apr 5]; 89:746–63.

12. Hu G, Hovav R, Grover CE, Faigenboim-Doron A, Kadmon N, Page JT, et al. Evolutionary Conservation and Divergence of Gene Coexpression Networks in Gossypium (Cotton) Seeds. Genome Biol Evol. Oxford University Press; 2017 [cited 2017 Apr 5];4:evw280.

13. Kawakatsu T, Huang SC, Jupe F, Sasaki E, Schmitz RJ, Urich MA, et al. Epigenomic Diversity in a Global Collection of Arabidopsis thaliana Accessions. Cell. 2016 [cited 2017 Apr 5];166:492–505.

14. Sircar S, Parekh N. Functional characterization of drought-responsive modules and genes in Oryza sativa: a network-based approach. Front Genet. Frontiers Media SA; 2015 [cited 2017 Apr 5];6:256.

15. Zinkgraf M, Liu L, Groover A, Filkov V. Identifying gene coexpression networks underlying the dynamic regulation of wood-forming tissues in Populus under diverse environmental conditions. New Phytol. 2017 [cited 2017 Apr 5];

16. Benfey PN, Mitchell-Olds T. From genotype to phenotype: systems biology meets natural variation. Science. NIH Public Access; 2008 [cited 2017 Apr 5];320:495–7.

17. Ogura T, Busch W. Genotypes, Networks, Phenotypes: Moving Toward Plant Systems Genetics. Annu Rev Cell Dev Biol. 2016 [cited 2017 Apr 5];32:103–26.

18. Dixit S, Grondin A, Lee C-R, Henry A, Olds T-M, Kumar A. Understanding rice adaptation to varying agro-ecosystems: trait interactions and quantitative trait loci. BMC Genet. BioMed Central; 2015 [cited 2017 Sep 28];16:86.

19. Langfelder P, Horvath S. WGCNA: an R package for weighted correlation network analysis. BMC Bioinformatics. 2008 [cited 2017 Apr 5];9:559.

20. Kauffmann A, Gentleman R, Huber W. arrayQualityMetrics-a bioconductor package for quality assessment of microarray data. Bioinformatics. 2009 [cited 2015 Sep 27];25:415–6.

21. Irizarry RA, Bolstad BM, Collin F, Cope LM, Hobbs B, Speed TP. Summaries of Affymetrix GeneChip probe level data. Nucleic Acids Res. 2003 [cited 2015 Jul 9];31:e15.

22. Mason MJ, Fan G, Plath K, Zhou Q, Horvath S. Signed weighted gene co-expression network analysis of transcriptional regulation in murine embryonic stem cells. BMC Genomics. 2009 [cited 2017 Jun 16];10:327.

23. Langfelder P, Zhang B, Horvath S. Defining clusters from a hierarchical cluster tree: the Dynamic Tree Cut package for R. Bioinformatics. 2008 [cited 2017 Apr 10];24:719–20.

24. Langfelder P, Luo R, Oldham MC, Horvath S. Is my network module preserved and reproducible? PLoS Comput Biol. 2011 [cited 2015 Apr 14];7:e1001057.

25. Du Z, Zhou X, Ling Y, Zhang Z, Su Z. agriGO: a GO analysis toolkit for the agricultural community. Nucleic Acids Res. 2010 [cited 2014 May 5];38:W64–70.

26. Szklarczyk D, Franceschini A, Wyder S, Forslund K, Heller D, Huerta-Cepas J, et al. STRING v10: protein-protein interaction networks, integrated over the tree of life. Nucleic Acids Res. 2015 [cited 2016 Jul 26];43:D447–52.

27. Lee T, Oh T, Yang S, Shin J, Hwang S, Kim CY, et al. RiceNet v2: an improved network prioritization server for rice genes. Nucleic Acids Res. Oxford University Press; 2015 [cited 2016 Jul 26];43:W122–7.

28. Lee T, Yang S, Kim E, Ko Y, Hwang S, Shin J, et al. AraNet v2: an improved database of co-functional gene networks for the study of Arabidopsis thaliana and 27 other nonmodel plant species. Nucleic Acids Res. 2014 [cited 2015 Jan 15];43:D996–1002.

29. Shannon P, Markiel A, Ozier O, Baliga NS, Wang JT, Ramage D, et al. Cytoscape: a software environment for integrated models of biomolecular interaction networks. Genome Res. 2003 [cited 2014 Jul 9];13:2498–504.

30. Pérez-Rodríguez P, Riaño-Pachón DM, Corrêa LGG, Rensing SA, Kersten B, Mueller-Roeber B. PlnTFDB: updated content and new features of the plant transcription factor database. Nucleic Acids Res. Oxford University Press; 2010 [cited 2017 Apr 10];38:D822–7.

31. Ozturk ZN, Talamé V, Deyholos M, Michalowski CB, Galbraith DW, Gozukirmizi N, et al. Monitoring large-scale changes in transcript abundance in drought-and salt-stressed barley. Plant Mol Biol. 2002 [cited 2017 Sep 24];48:551–73.

32. Hazen SP, Pathan MS, Sanchez A, Baxter I, Dunn M, Estes B, et al. Expression profiling of rice segregating for drought tolerance QTLs using a rice genome array. Funct Integr Genomics. 2005 [cited 2017 Sep 24];5:104–16.

33. Wang T, McFarlane HE, Persson S. The impact of abiotic factors on cellulose synthesis. J Exp Bot. Oxford University Press; 2016 [cited 2017 Sep 24];67:543–52.

34. Xu Z-Y, Kim SY, Hyeon DY, Kim DH, Dong T, Park Y, et al. The Arabidopsis NAC Transcription Factor ANAC096 Cooperates with bZIP-Type Transcription Factors in Dehydration and Osmotic Stress Responses. Plant Cell. 2013 [cited 2017 Sep 26];25:4708–24.

35. Nakashima K, Takasaki H, Mizoi J, Shinozaki K, Yamaguchi-Shinozaki K. NAC transcription factors in plant abiotic stress responses. Biochim Biophys Acta - Gene Regul Mech. 2012 [cited 2017 Sep 26];1819:97–103.

36. Jensen MK, Lindemose S, de Masi F, Reimer JJ, Nielsen M, Perera V, et al. ATAF1 transcription factor directly regulates abscisic acid biosynthetic gene NCED3 in Arabidopsis thaliana. FEBS Open Bio. 2013 [cited 2017 Sep 26];3:321–7.

37. Xie Z, Li D, Wang L, Sack FD, Grotewold E. Role of the stomatal development regulators FLP/MYB88 in abiotic stress responses. Plant J. 2010 [cited 2017 Sep 26];64:731–9.

38. Cominelli E, Galbiati M, Tonelli C. Transcription factors controlling stomatal movements and drought tolerance. Transcription. 2010 [cited 2017 Sep 26];1:41–5.

39. Yamaguchi-Shinozaki K, Shinozaki K. The plant hormone abscisic acid mediates the drought-induced expression but not the seed-specific expression of rd22, a gene responsive to dehydration stress in Arabidopsis thaliana. Mol Gen Genet. 1993 [cited 2017 Sep 26];238:17–25.

40. Lee S-j., Kang J-y., Park H-J, Kim MD, Bae MS, Choi H-i., et al. DREB2C Interacts with ABF2, a bZIP Protein Regulating Abscisic Acid-Responsive Gene Expression, and Its Overexpression Affects Abscisic Acid Sensitivity. PLANT Physiol. 2010 [cited 2018 Apr 16];153:716–27.

41. Jakoby M, Weisshaar B, Dröge-Laser W, Vicente-Carbajosa J, Tiedemann J, Kroj T, et al. bZIP transcription factors in Arabidopsis. Trends Plant Sci. 2002 [cited 2015 Jan 30];7:106–11.

42. Zhang C, Li C, Liu J, Lv Y, Yu C, Li H, et al. The OsABF1 transcription factor improves drought tolerance by activating the transcription of COR413-TM1 in rice. J Exp Bot. 2017 [cited 2017 Sep 17];171:2810–25.

43. Nuruzzaman M, Manimekalai R, Sharoni AM, Satoh K, Kondoh H, Ooka H, et al. Genome-wide analysis of NAC transcription factor family in rice. Gene. 2010 [cited 2017 Apr 4];465:30–44.

44. Gao F, Xiong A, Peng R, Jin X, Xu J, Zhu B, et al. OsNAC52, a rice NAC transcription factor, potentially responds to ABA and confers drought tolerance in transgenic plants. Plant Cell, Tissue Organ Cult. Springer Netherlands; 2010 [cited 2018 Apr 20];100:255–62.

45. Yanhui C, Xiaoyuan Y, Kun H, Meihua L, Jigang L, Zhaofeng G, et al. The MYB Transcription Factor Superfamily of Arabidopsis: Expression Analysis and Phylogenetic Comparison with the Rice MYB Family. Plant Mol Biol. 2006 [cited 2017 Apr 4];60:107–24.

46. Zhu N, Cheng S, Liu X, Du H, Dai M, Zhou D-X, et al. The R2R3-type MYB gene OsMYB91 has a function in coordinating plant growth and salt stress tolerance in rice. Plant Sci. 2015 [cited 2018 Feb 14];236:146–56.

47. Dietz K-J, Vogel MO, Viehhauser A. AP2/EREBP transcription factors are part of gene regulatory networks and integrate metabolic, hormonal and environmental signals in stress acclimation and retrograde signalling. Protoplasma. 2010 [cited 2017 Apr 4];245:3–14.

48. Sharoni AM, Nuruzzaman M, Satoh K, Shimizu T, Kondoh H, Sasaya T, et al. Gene Structures, Classification and Expression Models of the AP2/EREBP Transcription Factor Family in Rice. Plant Cell Physiol. Oxford University Press; 2011 [cited 2017 Apr 4];52:344–60.

49. Maruyama K, Takeda M, Kidokoro S, Yamada K, Sakuma Y, Urano K, et al. Metabolic Pathways Involved in Cold Acclimation Identified by Integrated Analysis of Metabolites and Transcripts Regulated by DREB1A and DREB2A. PLANT Physiol. 2009 [cited 2017 Apr 4];150:1972–80.

50. Qiu Y, Yu D. Over-expression of the stress-induced OsWRKY45 enhances disease resistance and drought tolerance in Arabidopsis. Environ Exp Bot. Elsevier; 2009 [cited 2017 Sep 27];65:35–47.

51. Raineri J, Wang S, Peleg Z, Blumwald E, Chan RL. The rice transcription factor OsWRKY47 is a positive regulator of the response to water deficit stress. Plant Mol Biol. 2015 [cited 2018 Feb 11];88:401–13.

52. Yonemaru J, Yamamoto T, Fukuoka S, Uga Y, Hori K, Yano M. Q-TARO: QTL Annotation Rice Online Database. Rice. Springer New York; 2010 [cited 2017 Apr 10];3:194–203.

53. Sairam RK, Tyagi A. Physiology and molecular biology of salinity stress tolerance in plants. Curr. Sci. Current Science Association; 2004 [cited 2018 Apr 6]. p. 407–21.

54. Usadel B, Poree F, Nagel A, Lohse M, Czedik-Eysenberg A, Stitt M. A guide to using MapMan to visualize and compare Omics data in plants: a case study in the crop species, Maize. Plant Cell Environ. 2009 [cited 2017 Jul 14];32:1211–29.

55. Childs KL, Davidson RM, Buell CR. Gene coexpression network analysis as a source of functional annotation for rice genes. El-Sayed NM, editor. PLoS One. Public Library of Science; 2011 [cited 2014 May 5];6:e22196.

56. Kanehisa M, Sato Y, Kawashima M, Furumichi M, Tanabe M. KEGG as a reference resource for gene and protein annotation. Nucleic Acids Res. 2016 [cited 2017 Jul 14];44:D457–62.

57. Reddy VS, Ali GS, Reddy ASN. Genes Encoding Calmodulin-binding Proteins in the Arabidopsis Genome. J Biol Chem. 2002 [cited 2017 Mar 20];277:9840–52.

58. Harkenrider M, Sharma R, De Vleesschauwer D, Tsao L, Zhang X, Chern M, et al. Overexpression of Rice Wall-Associated Kinase 25 (OsWAK25) Alters Resistance to Bacterial and Fungal Pathogens. Wang Z, editor. PLoS One. Public Library of Science; 2016 [cited 2017 Feb 23];11:e0147310.

59. Bayer RG, Stael S, Rocha AG, Mair A, Vothknecht UC, Teige M. Chloroplast-localized protein kinases: a step forward towards a complete inventory. J Exp Bot. Oxford University Press; 2012 [cited 2017 Feb 23];63:1713–23.

60. Zou Y, Liu X, Wang Q, Chen Y, Liu C, Qiu Y, et al. OsRPK1, a novel leucine-rich repeat receptor-like kinase, negatively regulates polar auxin transport and root development in rice. Biochim Biophys Acta. 2014 [cited 2017 Feb 23];1840:1676–85.

61. Gao L-L, Xue H-W. Global Analysis of Expression Profiles of Rice Receptor-Like Kinase Genes. Mol Plant. 2012;5:143–53.

62. Zong W, Tang N, Yang J, Peng L, Ma S, Xu Y, et al. Feedback Regulation of ABA Signaling and Biosynthesis by a bZIP Transcription Factor Targets Drought-Resistance-Related Genes. Plant Physiol. American Society of Plant Biologists; 2016 [cited 2016 Sep 2];171:2810–25. Available from: http://www.ncbi.nlm.nih.gov/pubmed/27325665

63. Hu H, You J, Fang Y, Zhu X, Qi Z, Xiong L. Characterization of transcription factor gene SNAC2 conferring cold and salt tolerance in rice. Plant Mol Biol. 2008 [cited 2017 Apr 10];67:169–81.

64. Nakashima K, Tran L-SP, Van Nguyen D, Fujita M, Maruyama K, Todaka D, et al. Functional analysis of a NAC-type transcription factor OsNAC6 involved in abiotic and biotic stress-responsive gene expression in rice. Plant J. Blackwell Publishing Ltd; 2007 [cited 2016 Jul 30];51:617–30.

65. Sun L, Huang L, Hong Y, Zhang H, Song F, Li D. Comprehensive Analysis Suggests Overlapping Expression of Rice ONAC Transcription Factors in Abiotic and Biotic Stress Responses. Int J Mol Sci. Multidisciplinary Digital Publishing Institute; 2015 [cited 2016 Jul 30];16:4306–26.

66. Hu H, You J, Fang Y, Zhu X, Qi Z, Xiong L. Characterization of transcription factor gene SNAC2 conferring cold and salt tolerance in rice. Plant Mol Biol. Springer Netherlands; 2008 [cited 2016 Jul 30];67:169–81.

67. Chae M-J, Lee J-S, Nam M-H, Cho K, Hong J-Y, Yi S-A, et al. A rice dehydration-inducible SNF1-related protein kinase 2 phosphorylates an abscisic acid responsive element-binding factor and associates with ABA signaling. Plant Mol Biol. Kluwer Academic Publishers; 2006 [cited 2016 Aug 11];63:151–69.

68. Kobayashi Y, Yamamoto S, Minami H, Kagaya Y, Hattori T. Differential activation of the rice sucrose nonfermenting1-related protein kinase2 family by hyperosmotic stress and abscisic acid. Plant Cell. American Society of Plant Biologists; 2004 [cited 2016 Aug 11];16:1163–77.

69. Nijhawan A, Jain M, Tyagi AK, Khurana JP. Genomic Survey and Gene Expression Analysis of the Basic Leucine Zipper Transcription Factor Family in Rice. PLANT Physiol. American Society of Plant Biologists; 2007 [cited 2016 Aug 19];146:333–50.

70. Hobo T, Kowyama Y, Hattori T. A bZIP factor, TRAB1, interacts with VP1 and mediates abscisic acid-induced transcription. Proc Natl Acad Sci U S A. National Academy of Sciences; 1999 [cited 2016 Aug 19];96:15348–53.

71. Kagaya Y, Hobo T, Murata M, Ban A, Hattori T. Abscisic acid-induced transcription is mediated by phosphorylation of an abscisic acid response element binding factor, TRAB1. Plant Cell. American Society of Plant Biologists; 2002 [cited 2016 Aug 19];14:3177–89.

72. Nijhawan A, Jain M, Tyagi AK, Khurana JP. Genomic survey and gene expression analysis of the basic leucine zipper transcription factor family in rice. Plant Physiol. American Society of Plant Biologists; 2008 [cited 2014 May 3];146:333–50.

73. Joo J, Lee YH, Song SI. Overexpression of the rice basic leucine zipper transcription factor OsbZIP12 confers drought tolerance to rice and makes seedlings hypersensitive to ABA. Plant Biotechnol Rep. Springer Japan; 2014 [cited 2016 Aug 19];8:431–41.

74. Izawa T, Foster R, Nakajima M, Shimamoto K, Chua NH. The rice bZIP transcriptional activator RITA-1 is highly expressed during seed development. Plant Cell. 1994 [cited 2016 Aug 19];6:1277–87.

75. Onodera Y, Suzuki A, Wu C-Y, Washida H, Takaiwa F. A Rice Functional Transcriptional Activator, RISBZ1, Responsible for Endosperm-specific Expression of Storage Protein Genes through GCN4 Motif. J Biol Chem. American Society for Biochemistry and Molecular Biology; 2001;276:14139–52.

76. Ma H-L, Xu X-H, Zhao X-Y, Liu H-J, Chen H. Impacts of drought stress on soluble carbohydrates and respiratory enzymes in fruit body of Auricularia auricula. Biotechnol Biotechnol Equip. Taylor & Francis; 2015 [cited 2016 Jul 31];29:10–4.

77. Chow C-N, Zheng H-Q, Wu N-Y, Chien C-H, Huang H-D, Lee T-Y, et al. PlantPAN 2.0: an update of plant promoter analysis navigator for reconstructing transcriptional regulatory networks in plants. Nucleic Acids Res. Oxford University Press; 2016 [cited 2018 Mar 23];44:D1154–60.

78. Li X-J, Yang M-F, Chen H, Qu L-Q, Chen F, Shen S-H. Abscisic acid pretreatment enhances salt tolerance of rice seedlings: proteomic evidence. Biochim Biophys Acta. 2010 [cited 2016 Sep 2];1804:929–40.

79. Hu W, Hu G, Han B. Genome-wide survey and expression profiling of heat shock proteins and heat shock factors revealed overlapped and stress specific response under abiotic stresses in rice. Plant Sci. 2009;176:583–90.

80. Lee B-H, Won S-H, Lee H-S, Miyao M, Chung W-I, Kim I-J, et al. Expression of the chloroplast-localized small heat shock protein by oxidative stress in rice. Gene. 2000;245:283–90.

81. Kelley WL. Molecular chaperones: How J domains turn on Hsp70s. Curr. Biol. 1999 [cited 2017 Apr 10]. p. R305–8.

82. Miot M, Reidy M, Doyle SM, Hoskins JR, Johnston DM, Genest O, et al. Species-specific collaboration of heat shock proteins (Hsp) 70 and 100 in thermotolerance and protein disaggregation. Proc Natl Acad Sci U S A. National Academy of Sciences; 2011 [cited 2017 Apr 10];108:6915–20.

83. Jacob P, Hirt H, Bendahmane A. The heat-shock protein/chaperone network and multiple stress resistance. Plant Biotechnol J. Wiley/Blackwell (10.1111); 2017 [cited 2018 Apr 14];15:405–14.

84. Singh A, Mittal D, Lavania D, Agarwal M, Mishra RC, Grover A. OsHsfA2c and OsHsfB4b are involved in the transcriptional regulation of cytoplasmic OsClpB (Hsp100) gene in rice (Oryza sativa L.). Cell Stress Chaperones. 2012 [cited 2017 Apr 1];17:243–54.

85. Le Gall H, Philippe F, Domon J-M, Gillet F, Pelloux J, Rayon C. Cell Wall Metabolism in Response to Abiotic Stress. Plants (Basel, Switzerland). Multidisciplinary Digital Publishing Institute (MDPI); 2015 [cited 2016 Aug 3];4:112–66.

86. Ricardi MM, González RM, Zhong S, Domínguez PG, Duffy T, Turjanski PG, et al. Genome-wide data (ChIP-seq) enabled identification of cell wall-related and aquaporin genes as targets of tomato ASR1, a drought stress-responsive transcription factor. BMC Plant Biol. BioMed Central; 2014 [cited 2016 Aug 3];14:29.

87. Kaul S, Sharma SS, Mehta IK. Free radical scavenging potential of L-proline: evidence from in vitro assays. Amino Acids. 2008 [cited 2017 Apr 10];34:315–20.

88. Yoshiba Y, Kiyosue T, Nakashima K, Yamaguchi-Shinozaki K, Shinozaki K. Regulation of levels of proline as an osmolyte in plants under water stress. Plant Cell Physiol. 1997 [cited 2017 Apr 10];38:1095–102.

89. Kavi Kishor PB, Hima Kumari P, Sunita MSL, Sreenivasulu N. Role of proline in cell wall synthesis and plant development and its implications in plant ontogeny. Front Plant Sci. Frontiers; 2015 [cited 2016 Aug 4];6:544.

90. Prabhu PR, Hudson AO, Prabhu PR, Hudson AO, Hudson A, O. Identification and Partial Characterization of an L-Tyrosine Aminotransferase (TAT) from Arabidopsis thaliana. Biochem Res Int. Hindawi Publishing Corporation; 2010 [cited 2016 Aug 4];2010:549572.

91. Mouradov A, Spangenberg G. Flavonoids: a metabolic network mediating plants adaptation to their real estate. Front Plant Sci. Frontiers Media SA; 2014 [cited 2017 Feb 23];5:620. Available from: http://www.ncbi.nlm.nih.gov/pubmed/25426130

92. Alvarez PJC (National AIC (Paraguay). D of G of BSC, Krzyzanowski FC, Mandarino JMG, Franca Neto JB. Relationship between soybean seed coat lignin content and resistance to mechanical damage. Seed Sci Technol. International Seed Testing Association; 1997;

93. Yoshimura K, Masuda A, Kuwano M, Yokota A, Akashi K. Programmed Proteome Response for Drought Avoidance/Tolerance in the Root of a C3 Xerophyte (Wild Watermelon) Under Water Deficits. Plant Cell Physiol. Oxford University Press; 2007 [cited 2017 Feb 23];49:226–41.

94. Li Y, Yuan F, Wen Z, Li Y, Wang F, Zhu T, et al. Genome-wide survey and expression analysis of the OSCA gene family in rice. BMC Plant Biol. 2015 [cited 2017 Mar 31];15:261.

95. Tiwari M, Sharma D, Singh M, Tripathi RD, Trivedi PK. Expression of OsMATE1 and OsMATE2 alters development, stress responses and pathogen susceptibility in Arabidopsis. Sci Rep. Nature Publishing Group; 2014 [cited 2017 Mar 31];4:1565–72.

96. Gaufichon L, Reisdorf-Cren M, Rothstein SJ, Chardon F, Suzuki A. Biological functions of asparagine synthetase in plants. Plant Sci. 2010 [cited 2017 Mar 31];179:141–53.

97. Araújo WL, Ishizaki K, Nunes-Nesi A, Larson TR, Tohge T, Krahnert I, et al. Identification of the 2-hydroxyglutarate and isovaleryl-CoA dehydrogenases as alternative electron donors linking lysine catabolism to the electron transport chain of Arabidopsis mitochondria. Plant Cell. 2010 [cited 2016 Aug 19];22:1549–63.

98. Araújo WL, Tohge T, Ishizaki K, Leaver CJ, Fernie AR. Protein degradation - an alternative respiratory substrate for stressed plants. Trends Plant Sci. 2011 [cited 2016 Aug 19];16:489–98.

99. Ishizaki K, Schauer N, Larson TR, Graham IA, Fernie AR, Leaver CJ. The mitochondrial electron transfer flavoprotein complex is essential for survival of Arabidopsis in extended darkness. Plant J. Blackwell Publishing Ltd; 2006 [cited 2016 Aug 19];47:751–60.

100. Binder S. Branched-Chain Amino Acid Metabolism in Arabidopsis thaliana. Arabidopsis Book. American Society of Plant Biologists; 2010 [cited 2016 Aug 17];8:e0137.

101. Finnegan PM, Soole KL, Umbach AL. Alternative Mitochondrial Electron Transport Proteins in Higher Plants. Springer Netherlands; 2004 [cited 2016 Aug 14]. p. 163–230.

102. Møller IM, Jensen PE, Hansson A. Oxidative modifications to cellular components in plants. Annu Rev Plant Biol. 2007 [cited 2016 Aug 14];58:459–81.

103. Wang J, Rajakulendran N, Amirsadeghi S, Vanlerberghe GC. Impact of mitochondrial alternative oxidase expression on the response of Nicotiana tabacum to cold temperature. Physiol Plant. 2011 [cited 2016 Aug 14];142:339–51.

104. Ma H, Song C, Borth W, Sether D, Melzer M, Hu J. Modified expression of alternative oxidase in transgenic tomato and petunia affects the level of tomato spotted wilt virus resistance. BMC Biotechnol. 2011 [cited 2016 Aug 14];11:96.

105. Vanlerberghe GC, Robson CA, Yip JYH. Induction of mitochondrial alternative oxidase in response to a cell signal pathway down-regulating the cytochrome pathway prevents programmed cell death. Plant Physiol. 2002 [cited 2016 Aug 14];129:1829–42.

106. Li C-R, Liang D-D, Li J, Duan Y-B, Li H, Yang Y-C, et al. Unravelling mitochondrial retrograde regulation in the abiotic stress induction of rice ALTERNATIVE OXIDASE 1 genes. Plant Cell Environ. Blackwell Publishing Ltd; 2013 [cited 2016 Aug 14];36:775–88.

107. Simova-Stoilova LP, Romero-Rodríguez MC, Sánchez-Lucas R, Navarro-Cerrillo RM, Medina-Aunon JA, Jorrín-Novo J V. 2-DE proteomics analysis of drought treated seedlings of Quercus ilex supports a root active strategy for metabolic adaptation in response to water shortage. Front Plant Sci. Frontiers Media SA; 2015 [cited 2016 Aug 14];6:627.

108. Alekseeva AA, Savin SS, Tishkov VI. NAD (+)-dependent Formate Dehydrogenase from Plants. Acta Naturae. Park Media; 2011 [cited 2016 Aug 14];3:38–54.

109. Qamar A, Mysore KS, Senthil-Kumar M. Role of proline and pyrroline-5-carboxylate metabolism in plant defense against invading pathogens. Front Plant Sci. Frontiers Media SA; 2015 [cited 2018 Feb 13];6:503.

110. Araújo WL, Nunes-Nesi A, Nikoloski Z, Sweetlove LJ, Fernie AR. Metabolic control and regulation of the tricarboxylic acid cycle in photosynthetic and heterotrophic plant tissues. Plant Cell Environ. Blackwell Publishing Ltd; 2012 [cited 2016 Aug 10];35:1–21.

111. Vasquez-Robinet C, Mane SP, Ulanov A V, Watkinson JI, Stromberg VK, De Koeyer D, et al. Physiological and molecular adaptations to drought in Andean potato genotypes. J Exp Bot. Oxford University Press; 2008 [cited 2016 Aug 10];59:2109–23.

112. Prasch CM, Sonnewald U. Simultaneous application of heat, drought, and virus to Arabidopsis plants reveals significant shifts in signaling networks. Plant Physiol. American Society of Plant Biologists; 2013 [cited 2016 Aug 10];162:1849–66.

113. Fatland BL, Nikolau BJ, Wurtele ES. Reverse Genetic Characterization of Cytosolic Acetyl-CoA Generation by ATP-Citrate Lyase in Arabidopsis. PLANT CELL ONLINE. 2005 [cited 2018 Feb 14];17:182–203.

114. Shi L, Tu BP. Acetyl-CoA and the regulation of metabolism: mechanisms and consequences. Curr Opin Cell Biol. NIH Public Access; 2015 [cited 2018 Feb 14];33:125–31.

115. Filichkin SA, Priest HD, Givan SA, Shen R, Bryant DW, Fox SE, et al. Genome-wide mapping of alternative splicing in Arabidopsis thaliana. Genome Res. Cold Spring Harbor Laboratory Press; 2010 [cited 2016 Aug 6];20:45–58.

116. Eichner J, Zeller G, Laubinger S, Rätsch G, Kim H, Klein R, et al. Support vector machines-based identification of alternative splicing in Arabidopsis thaliana from whole-genome tiling arrays. BMC Bioinformatics. BioMed Central; 2011 [cited 2016 Aug 6];12:55.

117. Ner-Gaon H, Halachmi R, Savaldi-Goldstein S, Rubin E, Ophir R, Fluhr R. Intron retention is a major phenomenon in alternative splicing in Arabidopsis. Plant J. Blackwell Science Ltd; 2004 [cited 2016 Aug 6];39:877–85.

118. Reed SI. Ratchets and clocks: the cell cycle, ubiquitylation and protein turnover. Nat Rev Mol Cell Biol. 2003 [cited 2016 Aug 7];4:855–64.

119. Hershko A. The ubiquitin system for protein degradation and some of its roles in the control of the cell division cycle. Cell Death Differ. 2005 [cited 2016 Aug 7];12:1191–7.

120. Kaur K, Gupta AK, Kaur N. Effect of water deficit on carbohydrate status and enzymes of carbohydrate metabolism in seedlings of wheat cultivars. Indian J Biochem Biophys. 2007 [cited 2016 Aug 17];44:223–30.

121. Zhao P, Liu P, Yuan G, Jia J, Li X, Qi D, et al. New Insights on Drought Stress Response by Global Investigation of Gene Expression Changes in Sheepgrass (Leymus chinensis). Front Plant Sci. Frontiers; 2016 [cited 2016 Aug 17];7:954.

122. Sharp RE. Interaction with ethylene: changing views on the role of abscisic acid in root and shoot growth responses to water stress. Plant, Cell Environ. Blackwell Science Ltd; 2002 [cited 2017 Apr 5];25:211–22.

123. Menkens AE, Schindler U, Cashmore AR. The G-box: a ubiquitous regulatory DNA element in plants bound by the GBF family of bZIP proteins. Trends Biochem Sci. 1995 [cited 2017 Apr 6];20:506–10.

124. Xie J, Tian J, Du Q, Chen J, Li Y, Yang X, et al. Association genetics and transcriptome analysis reveal a gibberellin-responsive pathway involved in regulating photosynthesis. J Exp Bot. Oxford University Press; 2016 [cited 2018 Mar 1];67:3325–38.

125. Schopfer P. Biomechanics of plant growth. Am J Bot. 2006 [cited 2017 Apr 6];93:1415–25.

126. Wang L, Guo K, Li Y, Tu Y, Hu H, Wang B, et al. Expression profiling and integrative analysis of the CESA/CSL superfamily in rice. BMC Plant Biol. BioMed Central; 2010 [cited 2016 Oct 4];10:282.

127. Noda S, Koshiba T, Hattori T, Yamaguchi M, Suzuki S, Umezawa T. The expression of a rice secondary wall-specific cellulose synthase gene, OsCesA7, is directly regulated by a rice transcription factor, OsMYB58/63. Planta. Springer Berlin Heidelberg; 2015 [cited 2016 Oct 4];242:589–600.

128. Vega-Sánchez ME, Verhertbruggen Y, Christensen U, Chen X, Sharma V, Varanasi P, et al. Loss of Cellulose synthase-like F6 function affects mixed-linkage glucan deposition, cell wall mechanical properties, and defense responses in vegetative tissues of rice. Plant Physiol. American Society of Plant Biologists; 2012 [cited 2016 Oct 4];159:56–69.

129. Gibeaut DM, Pauly M, Bacic A, Fincher GB. Changes in cell wall polysaccharides in developing barley (Hordeum vulgare) coleoptiles. Planta. 2005 [cited 2016 Oct 4];221:729–38.

131. Durner J, Shah J, Klessig DF. Salicylic acid and disease resistance in plants. Trends Plant Sci. 1997 [cited 2017 Apr 7];2:266–74.

132. Xu J, Audenaert K, Hofte M, De Vleesschauwer D. Abscisic Acid Promotes Susceptibility to the Rice Leaf Blight Pathogen Xanthomonas oryzae pv oryzae by Suppressing Salicylic Acid-Mediated Defenses. Yang C-H, editor. PLoS One. 2013 [cited 2017 Mar 24];8:e67413.

133. Jiang C-J, Shimono M, Sugano S, Kojima M, Yazawa K, Yoshida R, et al. Abscisic Acid Interacts Antagonistically with Salicylic Acid Signaling Pathway in Rice– Magnaporthe grisea Interaction. Mol Plant-Microbe Interact. 2010 [cited 2017 Mar 24];23:791–8.

134. Takatsuji H, Jiang C-J. Plant Hormone Crosstalks Under Biotic Stresses. Phytohormones A Wind to Metab Signal Biotechnol Appl. New York, NY: Springer New York; 2014 [cited 2017 Mar 24]. p. 323–50.

135. Dong X. NPR1, all things considered. Curr Opin Plant Biol. 2004 [cited 2017 Apr 7];7:547–52.

136. Cao H, Glazebrook J, Clarke JD, Volko S, Dong X. The Arabidopsis NPR1 gene that controls systemic acquired resistance encodes a novel protein containing ankyrin repeats. Cell. 1997 [cited 2017 Apr 7];88:57–63.

137. Jiang C-J, Shimono M, Sugano S, Kojima M, Yazawa K, Yoshida R, et al. Abscisic Acid Interacts Antagonistically with Salicylic Acid Signaling Pathway in Rice– Magnaporthe grisea Interaction. Mol Plant-Microbe Interact. 2010 [cited 2017 Apr 7];23:791–8.

138. Quilis J, Peñas G, Messeguer J, Brugidou C, Segundo BS. The Arabidopsis AtNPR1 Inversely Modulates Defense Responses Against Fungal, Bacterial, or Viral Pathogens While Conferring Hypersensitivity to Abiotic Stresses in Transgenic Rice. Mol Plant-Microbe Interact. 2008 [cited 2017 Mar 24];21:1215–31.

139. Wu T, Tian Z, Liu J, Yao C, Xie C. A Novel Ankyrin Repeat-rich Gene in Potato, Star, Involved in Response to Late Blight. Biochem Genet. 2009 [cited 2016 Nov 11];47:439–50.

140. Sacco MA, Mansoor S, Moffett P. A RanGAP protein physically interacts with the NB-LRR protein Rx, and is required for Rx-mediated viral resistance. Plant J. 2007 [cited 2016 Nov 11];52:82–93.

141. Jacques A, Ghannam A, Erhardt M, de Ruffray P, Baillieul F, Kauffmann S. NtLRP1, a Tobacco Leucine-Rich Repeat Gene with a Possible Role as a Modulator of the Hypersensitive Response. Mol Plant-Microbe Interact. 2006 [cited 2016 Nov 11];19:747–57.

142. Mou S, Liu Z, Guan D, Qiu A, Lai Y, He S. Functional analysis and expressional characterization of rice ankyrin repeat-containing protein, OsPIANK1, in basal defense against Magnaporthe oryzae attack. PLoS One. Public Library of Science; 2013 [cited 2016 Nov 11];8:e59699.

143. Kwak S-H. Positional Signaling Mediated by a Receptor-like Kinase in Arabidopsis. Science (80-). 2005 [cited 2016 Nov 8];307:1111–3.

144. Kwak S-H, Schiefelbein J. The role of the SCRAMBLED receptor-like kinase in patterning the Arabidopsis root epidermis. Dev Biol. 2007 [cited 2016 Nov 8];302:118–31.

145. Chevalier D, Batoux M, Fulton L, Pfister K, Yadav RK, Schellenberg M, et al. STRUBBELIG defines a receptor kinase-mediated signaling pathway regulating organ development in Arabidopsis. Proc Natl Acad Sci. 2005 [cited 2016 Nov 8];102:9074–9.

146. Chern M, Fitzgerald HA, Canlas PE, Navarre DA, Ronald PC. Overexpression of a Rice NPR1 Homolog Leads to Constitutive Activation of Defense Response and Hypersensitivity to Light. Mol Plant-Microbe Interact. 2005 [cited 2016 Nov 11];18:511–20.

147. Tang J, Zhu X, Wang Y, Liu L, Xu B, Li F, et al. Semi-dominant mutations in the CC-NB-LRR-type R gene, NLS1, lead to constitutive activation of defense responses in rice. Plant J. 2011 [cited 2016 Nov 11];66:996–1007.

148. Feng J-X, Cao L, Li J, Duan C-J, Luo X-M, Le N, et al. Involvement of OsNPR1/NH1 in rice basal resistance to blast fungus Magnaporthe oryzae. Eur J Plant Pathol. Springer Netherlands; 2011 [cited 2016 Nov 11];131:221–35.

149. van Ooijen G, Mayr G, Kasiem MMA, Albrecht M, Cornelissen BJC, Takken FLW. Structure-function analysis of the NB-ARC domain of plant disease resistance proteins. J Exp Bot. Oxford University Press; 2008 [cited 2017 Feb 27];59:1383–97.

150. Liu Q, Yang J, Zhang S, Zhao J, Feng A, Yang T, et al. OsGF14e positively regulates panicle blast resistance in rice. Biochem. Biophys. Res. Commun. 2016.

151. Liu Q, Yang J, Zhang S, Zhao J, Feng A, Yang T, et al. OsGF14b Positively Regulates Panicle Blast Resistance but Negatively Regulates Leaf Blast Resistance in Rice. Mol Plant-Microbe Interact. 2016 [cited 2017 Feb 27];29:46–56.

152. Pignocchi C, Doonan JH. Interaction of a 14-3-3 protein with the plant microtubule-associated protein EDE1. Ann Bot. Oxford University Press; 2011 [cited 2017 Feb 27];107:1103–9.

153. Kressler D, Hurt E, Baβler J. Driving ribosome assembly. Biochim Biophys Acta - Mol Cell Res. Elsevier; 2010 [cited 2018 Mar 3];1803:673–83.

154. Henras AK, Soudet J, Gérus M, Lebaron S, Caizergues-Ferrer M, Mougin A, et al. The post-transcriptional steps of eukaryotic ribosome biogenesis. Cell Mol Life Sci. 2008 [cited 2018 Mar 3];65:2334–59.

155. Ye H, Du H, Tang N, Li X, Xiong L. Identification and expression profiling analysis of TIFY family genes involved in stress and phytohormone responses in rice. Plant Mol Biol. 2009 [cited 2018 Mar 4];71:291–305.

156. Parenicová L, de Folter S, Kieffer M, Horner DS, Favalli C, Busscher J, et al. Molecular and phylogenetic analyses of the complete MADS-box transcription factor family in Arabidopsis: new openings to the MADS world. Plant Cell. American Society of Plant Biologists; 2003 [cited 2018 Mar 4];15:1538–51.

157. Ledger S, Strayer C, Ashton F, Kay SA, Putterill J. Analysis of the function of two circadian-regulated CONSTANS-LIKE genes. Plant J. 2001 [cited 2018 Mar 4];26:15–22.

158. Putterill J, Robson F, Lee K, Simon R, Coupland G. The CONSTANS gene of Arabidopsis promotes flowering and encodes a protein showing similarities to zinc finger transcription factors. Cell. 1995 [cited 2018 Mar 4];80:847–57.

159. Du L, Jiao F, Chu J, Jin G, Chen M, Wu P. The two-component signal system in rice (Oryza sativa L.): A genome-wide study of cytokinin signal perception and transduction. Genomics. Academic Press; 2007 [cited 2018 Mar 4];89:697–707.

160. Jain M, Tyagi AK, Khurana JP. Molecular characterization and differential expression of cytokinin-responsive type-A response regulators in rice (Oryza sativa). BMC Plant Biol. BioMed Central; 2006 [cited 2018 Mar 4];6:1.

161. Turner JG, Ellis C, Devoto A. The jasmonate signal pathway. Plant Cell. American Society of Plant Biologists; 2002 [cited 2017 Apr 10];14 Suppl:S153–64.

162. Wakuta S, Suzuki E, Saburi W, Matsuura H, Nabeta K, Imai R, et al. OsJAR1 and OsJAR2 are jasmonyl-l-isoleucine synthases involved in wound-and pathogen-induced jasmonic acid signalling. Biochem Biophys Res Commun. 2011 [cited 2017 Feb 27];409:634–9.

163. Haga K, Iino M. Phytochrome-mediated transcriptional up-regulation of ALLENE OXIDE SYNTHASE in rice seedlings. Plant Cell Physiol. 2004 [cited 2017 Mar 28];45:119–28.

164. Tani T, Sobajima H, Okada K, Chujo T, Arimura S, Tsutsumi N, et al. Identification of the OsOPR7 gene encoding 12-oxophytodienoate reductase involved in the biosynthesis of jasmonic acid in rice. Planta. 2008 [cited 2017 Mar 28];227:517–26.

165. Chehab EW, Raman G, Walley JW, Perea J V, Banu G, Theg S, et al. Rice HYDROPEROXIDE LYASES with unique expression patterns generate distinct aldehyde signatures in Arabidopsis. Plant Physiol. 2006 [cited 2017 Mar 28];141:121–34.

166. Hayes JD, Flanagan JU, Jowsey IR. GLUTATHIONE TRANSFERASES. Annu Rev Pharmacol Toxicol. 2005 [cited 2018 Mar 22];45:51–88.

167. Jain M, Ghanashyam C, Bhattacharjee A. Comprehensive expression analysis suggests overlapping and specific roles of rice glutathione S-transferase genes during development and stress responses. BMC Genomics. BioMed Central; 2010 [cited 2018 Mar 22];11:73.

168. Wingler A, Lea PJ, Quick WP, Leegood RC. Photorespiration: metabolic pathways and their role in stress protection. Philos Trans R Soc Lond B Biol Sci. The Royal Society; 2000 [cited 2017 Mar 28];355:1517–29.

169. Yin H, Zhang X, Liu J, Wang Y, He J, Yang T, et al. Epigenetic regulation, somatic homologous recombination, and abscisic acid signaling are influenced by DNA polymerase epsilon mutation in Arabidopsis. Plant Cell. 2009 [cited 2017 Mar 30];21:386–402.

170. He J, Duan Y, Hua D, Fan G, Wang L, Liu Y, et al. DEXH box RNA helicase-mediated mitochondrial reactive oxygen species production in Arabidopsis mediates crosstalk between abscisic acid and auxin signaling. Plant Cell. 2012 [cited 2017 Mar 30];24:1815–33.

171. Wang L, Hua D, He J, Duan Y, Chen Z, Hong X, et al. Auxin Response Factor2 (ARF2) and its regulated homeodomain gene HB33 mediate abscisic acid response in Arabidopsis. Qu L-J, editor. PLoS Genet. 2011 [cited 2017 Mar 30];7:e1002172.

172. Titiz O, Tambasco-Studart M, Warzych E, Apel K, Amrhein N, Laloi C, et al. PDX1 is essential for vitamin B6 biosynthesis, development and stress tolerance in Arabidopsis. Plant J. 2006 [cited 2018 Mar 18];48:933–46.

173. Wagner S, Bernhardt A, Leuendorf JE, Drewke C, Lytovchenko A, Mujahed N, et al. Analysis of the Arabidopsis rsr4-1/pdx1-3 Mutant Reveals the Critical Function of the PDX1 Protein Family in Metabolism, Development, and Vitamin B6 Biosynthesis. PLANT CELL ONLINE. 2006 [cited 2018 Mar 18];18:1722–35.

174. Bassel GW, Lan H, Glaab E, Gibbs DJ, Gerjets T, Krasnogor N, et al. Genome-wide network model capturing seed germination reveals coordinated regulation of plant cellular phase transitions. Proc Natl Acad Sci. 2011 [cited 2017 Apr 10];108:9709–14.

175. Massonnet M, Morales-Cruz A, Figueroa-Balderas R, Lawrence DP, Baumgartner K, Cantu D. Condition-dependent co-regulation of genomic clusters of virulence factors in the grapevine trunk pathogen Neofusicoccum parvum. Mol Plant Pathol. 2016 [cited 2017 Apr 10];

176. Schaefer RJ, Briskine R, Springer NM, Myers CL, Morris Q. Discovering Functional Modules across Diverse Maize Transcriptomes Using COB, the Co-Expression Browser. Börnke F, editor. PLoS One. Public Library of Science; 2014 [cited 2017 Apr 10];9:e99193.

177. Shuqin Zhang S, Hongyu Zhao H, Ng MK. Functional Module Analysis for Gene Coexpression Networks with Network Integration. IEEE/ACM Trans Comput Biol Bioinforma. 2015 [cited 2017 Apr 10];12:1146–60.

178. Ruprecht C, Vaid N, Proost S, Persson S, Mutwil M. Beyond Genomics: Studying Evolution with Gene Coexpression Networks. Trends Plant Sci. 2017 [cited 2017 Apr 10];22:298–307.

179. Tzfadia O, Diels T, De Meyer S, Vandepoele K, Aharoni A, Van de Peer Y. CoExpNetViz: Comparative Co-Expression Networks Construction and Visualization Tool. Front Plant Sci. Frontiers Media SA; 2015 [cited 2017 Apr 10];6:1194.

180. Yokotani N, Ichikawa T, Kondou Y, Matsui M, Hirochika H, Iwabuchi M, et al. Expression of rice heat stress transcription factor OsHsfA2e enhances tolerance to environmental stresses in transgenic Arabidopsis. Planta. 2008 [cited 2017 Sep 28];227:957–67.

181. Braunstein Ae. [Principal ways of assimilation & dissimilation of nitrogen in animals]. Adv Enzymol Relat Subj Biochem. 1957 [cited 2016 Sep 12];19:335–89.

